# The metabolic program of inflammatory eosinophils accounts for chronic parasite-induced skin disease

**DOI:** 10.1101/2025.02.28.640104

**Authors:** David Barinberg, Heidi Sebald, Tobias Gold, Baplu Rai, Daniel Radtke, Dominik Lerm, David Voehringer, Jonathan Jantsch, Stefan Wirtz, Alina Ulezko Antonova, Marco Colonna, Christian Bogdan, Ulrike Schleicher

## Abstract

Eosinophils exert antimicrobial, cytotoxic and immunoregulatory effects, but their function in cutaneous tissue still remains poorly understood. Here, we used a mouse model of chronic cutaneous leishmaniasis caused by the protozoan parasite *Leishmania (L.) mexicana* to investigate the function and transcriptomic signature of eosinophils in the skin. In C57BL/6 wild-type mice, *L. mexicana* infection induced local and systemic eosinophilia that was dependent on type 2 innate lymphoid cells and interleukin-5. Genetic and pharmacological depletion of eosinophils led to complete clinical resolution of disease, which was accompanied by a more pronounced Th1 and M1-like macrophage response. Bioinformatic analyses revealed a novel inflammatory and tissue-specific transcriptional trajectory in skin-infiltrating eosinophils. Skin-imprinted eosinophils strongly expressed the high-affinity glucose transporter 3 (*Slc2a3*), deprived the environment of glucose and directly impeded the function of Th1 cells. Together, our results demonstrate that disease progression and chronicity of *L. mexicana* infection is caused by inflammatory eosinophils and linked to their metabolic program.

**Short Summary:** The authors describe that depletion of eosinophils prevents chronic cutaneous disease caused by *Leishmania mexicana*. They identify a novel, tissue-specific transcriptomic profile of inflammatory skin eosinophils and demonstrate that skin-imprinted eosinophils show strong glucose uptake and directly repress Th1 responses.

**GRAPHICAL ABSTRACT:** 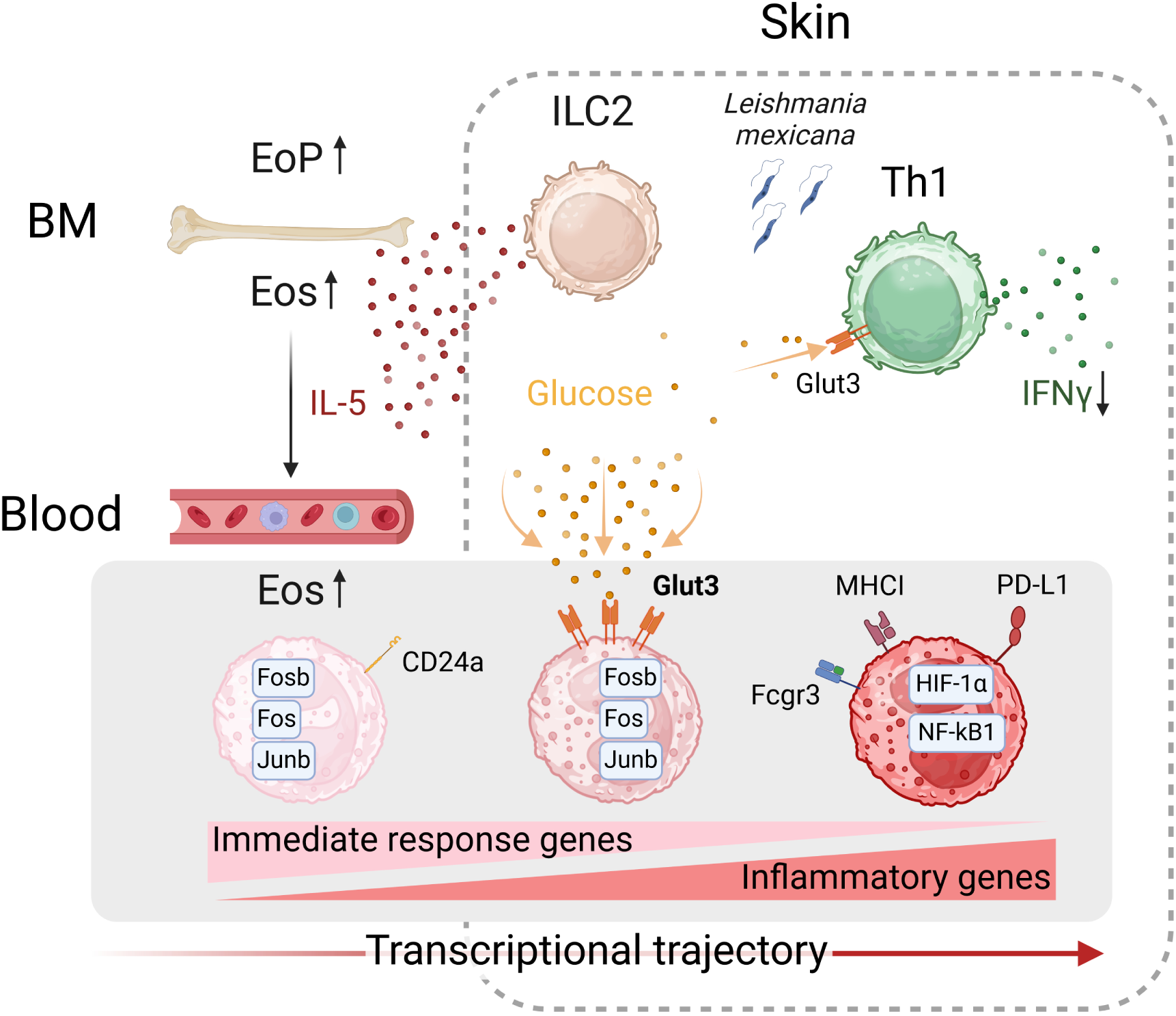

**KEY POINTS:**

- Eosinophil accumulation precedes the development of chronic cutaneous leishmaniasis
- Eosinophil depletion or IL-5 neutralization lead to clinical cure of the disease
- *L. mexicana* infection elicits a unique transcriptomic signature of skin eosinophils
- Skin eosinophils show a marked uptake of glucose and directly repress Th1 responses

## INTRODUCTION

Eosinophils are a distinct subset of granulocytes derived from bone marrow (BM) myeloid progenitors. They accumulate in various tissues, e.g., in the thymus (Muller, 1977), uterus (Gouon-Evans, 2001), lungs (Mesnil et al., 2016), adipose tissue (Wu et al., 2011), mammary gland (Gouon-Evans et al., 2000) and gastrointestinal tract (Chu et al., 2014), whereas the peripheral blood contains only few eosinophils under homeostatic conditions. For a long time, eosinophils have been viewed as typical effector cells during the resolution phase of inflammations (Gleich, 2013; Isobe et al., 2012) or during diseases characterized by type 2 immune reactions, such as allergies and asthma (Rosenberg et al., 2007) and helminth infections (Huang and Appleton, 2016). Over the last decade, our knowledge of the function of eosinophils has significantly increased (Arnold and Munitz, 2024). Eosinophils were recognized to contribute to the control of bacterial infections (Bohrer et al., 2021), to support lymphocytes in their protective activity against cancer (Blomberg et al., 2023; Carretero et al., 2015; Grisaru-Tal et al., 2022; Tepper et al., 1992) and to be critical for the immune homeostasis in a variety of organs (Brigger et al., 2020; Gurtner et al., 2023; Ignacio et al., 2022; Mesnil et al., 2016; Wu et al., 2011). In contrast, the role of eosinophils in chronic cutaneous infections remains poorly understood and requires further investigation, considering the barrier and antimicrobial function of the skin (Coates et al., 2018).

While the skin is typically devoid of eosinophils under physiological conditions, eosinophil infiltrations are characteristic for a broad spectrum of skin diseases (Radonjic-Hoesli et al., 2021). These include a wide range of inflammatory dermatoses (e.g., allergic drug eruption, urticaria, atopic dermatitis, eczema, allergic contact dermatitis, and other conditions with prurigo) and malignancies that may affect people at all ages (Heymann, 2006; Leiferman and Peters, 2018; Long et al., 2016; Montgomery et al., 2013). Despite their high prevalence, the exact pathogenic mechanisms of eosinophils remain obscure for most eosinophilic dermatoses (Radonjic-Hoesli et al., 2021).

Cutaneous leishmaniasis (CL) is a vector-borne disease caused by different species of the intracellular protozoan parasite *Leishmania* (Bogdan et al., 2019; Scott and Novais, 2016). Parasite and disease control is critically dependent on interferon (IFN)γ-producing type 1 T helper (Th1) cells and macrophages expressing type 2 nitric oxide synthase (NOS2), whereas non-healing, chronic CL has been linked to the expansion of interleukin (IL)-4-secreting Th2 cells, IL-10-producing CD4^+^ T cells and the expression of arginase 1 (ARG1) (Bogdan et al., 2024; Buxbaum, 2015; Padigel et al., 2003; Sacks and Noben-Trauth, 2002; Schleicher et al., 2016). Information on the role of eosinophils in CL has been scarce. Immunohistology detected eosinophils in the vicinity of macrophages in *Leishmania*-infected skin lesions of mice or man (Beil et al., 1992; Grimaldi et al., 1984; McElrath et al., 1987; Pompeu et al., 1991)*. In vitro* studies suggested that eosinophils participate in the killing of *Leishmania* parasites: either directly, via the release of granules (with antimicrobial peptides), reactive oxygen species (ROS) and nitric oxide (NO), or the formation of eosinophil extracellular DNA traps (termed EETosis) (Oliveira et al., 1998; Pimenta et al., 1987; Salaiza-Suazo et al., 2024; Watanabe et al., 2004); or indirectly via the stimulation of NO-dependent antileishmanial activity of macrophages by eosinophil-derived prostaglandin D2 (da Silva Marques et al., 2021). In line with these protective *in vitro* effects, eosinophil-deficient BALB/c dblGATA-1 mice developed a much more severe course of *L. amazonensis* infection as compared to WT controls (Almeida et al., 2024). Finally, recent investigations in a C57BL/6 mouse model of rapidly progressive CL due to infection with a special *L. major* strain (Seidman) revealed a regulatory function of eosinophils. Eosinophils, which were recruited in response to eotaxin 2 (CCL24)- and thymic stromal lymphopoietin (TSLP)-secreting tissue-resident macrophages (TRM) and IL-5-releasing type 2 innate lymphoid cells (ILC2s), produced IL-4 that helped maintaining dermal TRM (*Mrc1*^high^ *Cd163*^+^) as a source of CCL24 and niche for parasite replication. Importantly, however, deletion of eosinophil-derived IL-4, ILC2-derived IL-5 or TSLP only ameliorated the skin lesion development, but did not prevent progression and chronicity of disease (Lee et al., 2020; Lee et al., 2023).

In the present study, we carried out a comprehensive analysis of the function of eosinophils in chronic persistent CL, combining *in vivo* experiments with multicolour flow cytometry, RT-qPCR and transcriptomic profiling of skin-infiltrating and circulating eosinophils by nanowell-based single-cell RNA sequencing (scRNA-seq). For infection, we chose *L. mexicana*, which causes chronic skin lesions without spontaneous healing not only in Th2-biased BALB/c and Th1-prone C57BL/6 mice (Aguilar Torrentera et al., 2002; Rosas et al., 2005), but also frequently in humans (Salaiza-Suazo et al., 2024). In C57BL/6 mice, *L. mexicana* induced cutaneous and systemic eosinophilia that was dependent on IL-5 and ILC2s. Remarkably, depletion of eosinophils led to a self-healing clinical course of infection that was accompanied by an altered composition of the monocyte/macrophage and the T/NK-cell compartments. Mechanistically, we observed that skin lesion-derived eosinophils rapidly took up glucose and suppressed the function of Th1 cells. Computational analyses revealed a previously uncharacterized inflammatory transcriptional trajectory in tissue-infiltrating eosinophils. Together, our results point towards a hitherto unknown, highly inflammatory eosinophil subset that plays a pivotal role in the progression of *L. mexicana* infection.

## RESULTS

### *L. mexicana* infection induces an IL-5-dependent local and systemic eosinophilia

In *L. mexicana*-infected C57BL/6 mice, eosinophilic granulocytes (SSC^high^ CD11b^+^ SiglecF^+^ cells) accumulated at the skin site of infection as early as day 6 post infection (p.i.) (**Figure 1A**, left; **Figure S1A**). The initial influx of eosinophils peaked at day 25 to 30 p.i., when around 70% of all viable cells were eosinophils, and was followed by a rapid decline thereafter, whereas the size of the skin lesions continuously increased during the observation period (**Figure 1A**, right). As IL-5 is critical for recruitment, expansion and activation of eosinophils in mice (Dougan et al., 2019; Jacobsen et al., 2021; Jorssen et al., 2024), we analysed its expression during the course of infection. On day 14 p.i., we noted an upregulation of *Il5* mRNA in the skin (**Figure 1B**) and a maximum level of IL-5 protein in the serum as detected by bead-based multiplex ELISA (**Figure 1C**) and IL-5-specific ELISA (**Figure 1D**). Since IL-5 is associated with systemic eosinophilia (Takatsu and Nakajima, 2008), we also investigated bone marrow (BM) and peripheral blood. In accordance with the IL-5 serum data, flow cytometry revealed a peak of mature eosinophils (Eos; CD45^+^ CD11b^+^ SiglecF^+^) and progenitor eosinophils (EoPs; CD11b^+^ CD34^+^ CD125^+^ SiglecF^+^) in the BM and of eosinophils in the blood (**Figure 1E, Figure S1B**). To test, whether IL-5 drives the expansion of eosinophils after *L. mexicana* infection in C57BL/6 mice, we applied a neutralizing anti-IL-5 antibody on day 5 and day 12 p.i. (**Figure S1C**) and found that the numbers of eosinophils in BM, blood and skin lesions were strongly reduced (**Figure 1F**). Thus, an infection with *L. mexicana* causes an IL-5-dependent local and systemic eosinophilia.

**Figure 1:**
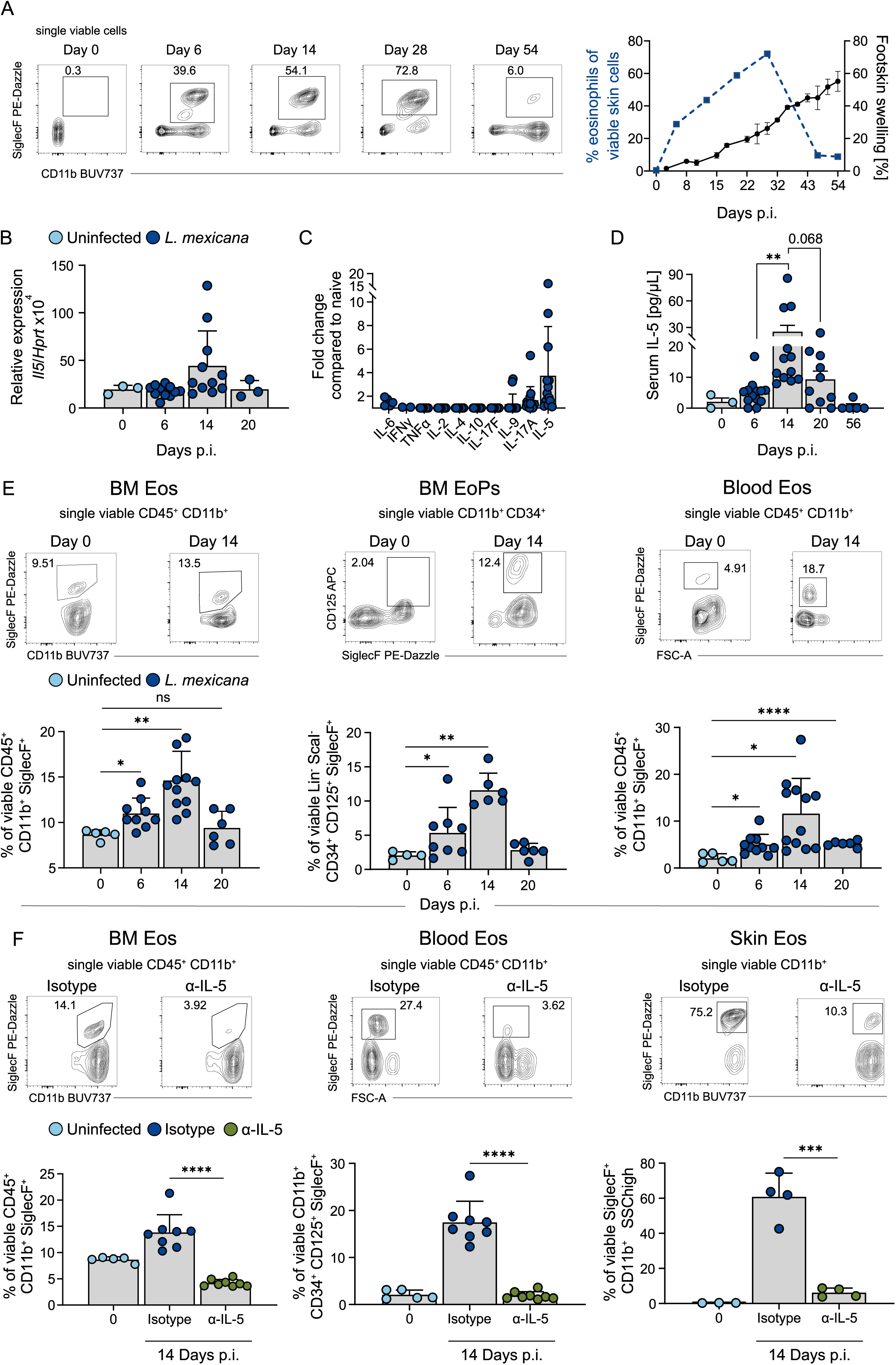
*L. mexicana* infection induces an IL-5-dependent local and systemic eosinophilia **A** Left, representative flow cytometric analysis of isolated C57BL/6 foot skin cells gated on CD11b^+^ SiglecF^+^ eosinophils on the respective days after *L. mexicana* infection. Right, representative clinical course of C57BL/6 mice infected with *L. mexicana* and influx of SSC^high^ CD11b^+^ SiglecF^+^ eosinophils as measured by flow cytometry (n = 4-6 mice per group, 4 independent experiments). **B** Quantification of *Il5* mRNA expression by qRT-PCR in *L. mexicana*-induced skin lesions of C57BL/6 mice (n = 3-7 mice per group, 1-3 independent experiments). **C** Bead-based multiplex ELISA of serum cytokines from C57BL/6 mice 14 days p.i. with *L. mexicana* (n = 4-14 mice per group, 3-4 independent experiments). **D** Serum IL-5 levels measured by ELISA in C57BL/6 mice infected with *L. mexicana* (n = 3-12 mice per group, 2-4 independent experiments). **E** Top, representative flow cytometric analysis of isolated cells from bone marrow and blood of naïve C57BL/6 mice or infected with *L. mexicana.* Bottom, quantification of the respective population and organ (n = 4-9 mice per group, 1-3 independent experiments). **F** C57BL/6 mice were treated with 500 µg of either isotype control or anti-IL-5 antibody on days 5 and 12 p.i.. Top, representative flow cytometric analysis of isolated cells from bone marrow, blood and skin lesions of isotype-treated or anti-IL-5-treated mice. Bottom, quantification of the respective population and organ. (n = 3-8 mice per group, 2 independent experiments; skin data points were derived from samples pooled from two mice). **A-F** *p ≤ 0.05; **p ≤ 0.01; ****p ≤ 0.0001, two-tailed Mann-Whitney U test. Data are mean ± SEM **(A)** or mean ± s.d. **(B-F).**

### ILC2s are essential for the *L. mexicana-*induced local and systemic eosinophilia

IL-5 is produced by Th2, ILC2s, mast cells and bone marrow stromal cells, but also by eosinophils themselves (Dougan et al., 2019; Hogan et al., 2008). As an accumulation of eosinophils in *L. mexicana*-infected skin was already visible a few days after infection, we first studied the role of ILC2s by infecting RorαΔTek (KO) and Rorα fl/fl (WT) mice. The KO mice, which are deficient for the transcription factor RAR-related orphan receptor alpha (*Rora)* in hematopoietic and endothelial cells, exhibit an almost complete absence of steady-state pulmonary IL-5- and IL-13-producing ILC2s (Kindermann et al., 2020; Knipfer et al., 2019). *L. mexicana*-infected RorαΔTek mice neither showed an increase in serum IL-5 (**Figure 2A**) nor the usual expansion of eosinophils (CD45^+^ CD11b^+^ SiglecF^+^) and eosinophil progenitors (EoPs; CD11b^+^ CD34^+^ CD125^+^ SiglecF^+^) in BM or of total eosinophils in blood and infected skin (**Figure 2B** and **C**). In contrast, the expansion of eosinophils in BM and blood at day 6 p.i. was maintained in *L. mexicana*-infected Rag1^-/-^ mice, which lack T and B lymphocytes but retain ILC2s (**Figure S1D**). Notably, by day 14 p.i., the magnitude of eosinophil expansion in the BM of Rag1^-/-^ mice was diminished compared to C57BL/6 WT controls, suggesting a partial contribution of the adaptive immune system to eosinophilia during later stages of infection (**Figure S1D**). Together, these findings demonstrate that ILC2s are required for the IL-5-dependent systemic and local eosinophilia early after *L. mexicana*-infection.

**Figure 2:**
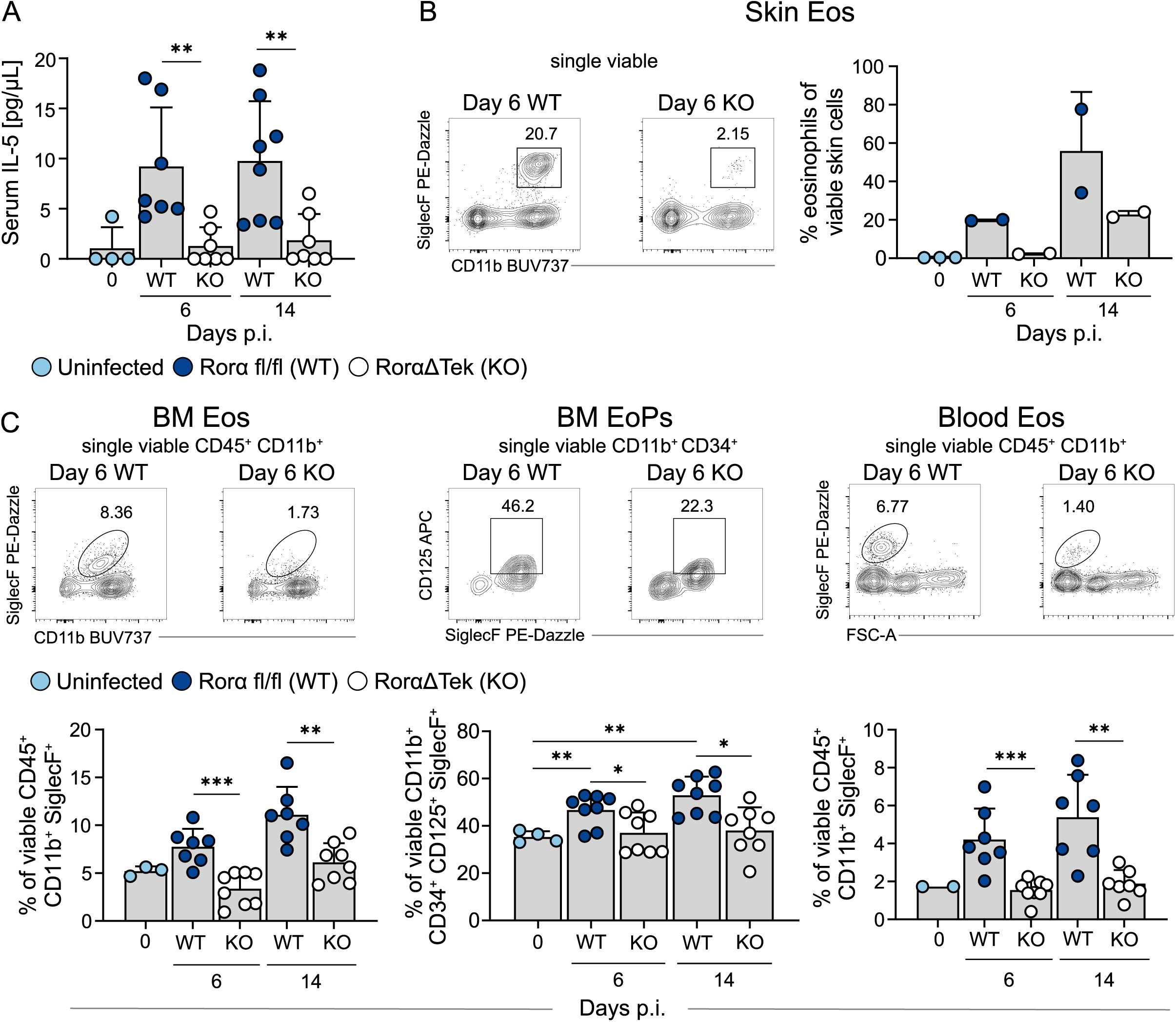
ILC2s are essential for the *L. mexicana-*induced local and systemic eosinophilia **A** Serum IL-5 levels measured by ELISA in Rorα fl/fl (WT) and RorαΔTek (KO) mice infected with *L. mexicana* (n = 4-8 mice per group, 2 independent experiments). **B** Left, representative flow cytometric analysis of isolated skin cells from WT and KO mice infected with *L. mexicana*. Right, quantification of the respective population (n = 2-3 mice per group, one experiment; skin data points were derived from samples pooled from two mice). **C** Top, representative flow cytometric analysis of isolated cells from bone marrow and blood of WT and KO mice infected with *L. mexicana*. Bottom, quantification of the respective population and organ (n = 2-8 mice per group, 2 independent experiments). *p ≤ 0.05; **p ≤ 0.01; ***p ≤ 0.005, two-tailed Mann-Whitney U test. Data are mean ± s.d.

### Lack of eosinophils or inhibition of eosinophil expansion prevents chronicity of *L. mexicana* infection

To elucidate the functional relevance of eosinophils during *L. mexicana* infection, we studied the course of *L. mexicana* infection in ΔdblGATA-1 4get C57BL/6 (dblGATA-1) mice, which lack eosinophils due to a targeted deletion of the GATA-1 binding site in the GATA-1 promoter (Yu et al., 2002) and show EGFP reporter expression in *Il4*-expressing cells (Mohrs et al., 2001). Remarkably, dblGATA-1 mice allowed for resolution of *L. mexicana* infection, whereas 4get C57BL/6 (WT) controls developed the expected chronic disease (**Figure 3A**). Cure of disease was associated with a substantial decrease in parasite burden in the foot skin, draining lymph node (dLN) and spleen (**Figure 3B**).

**Figure 3:**
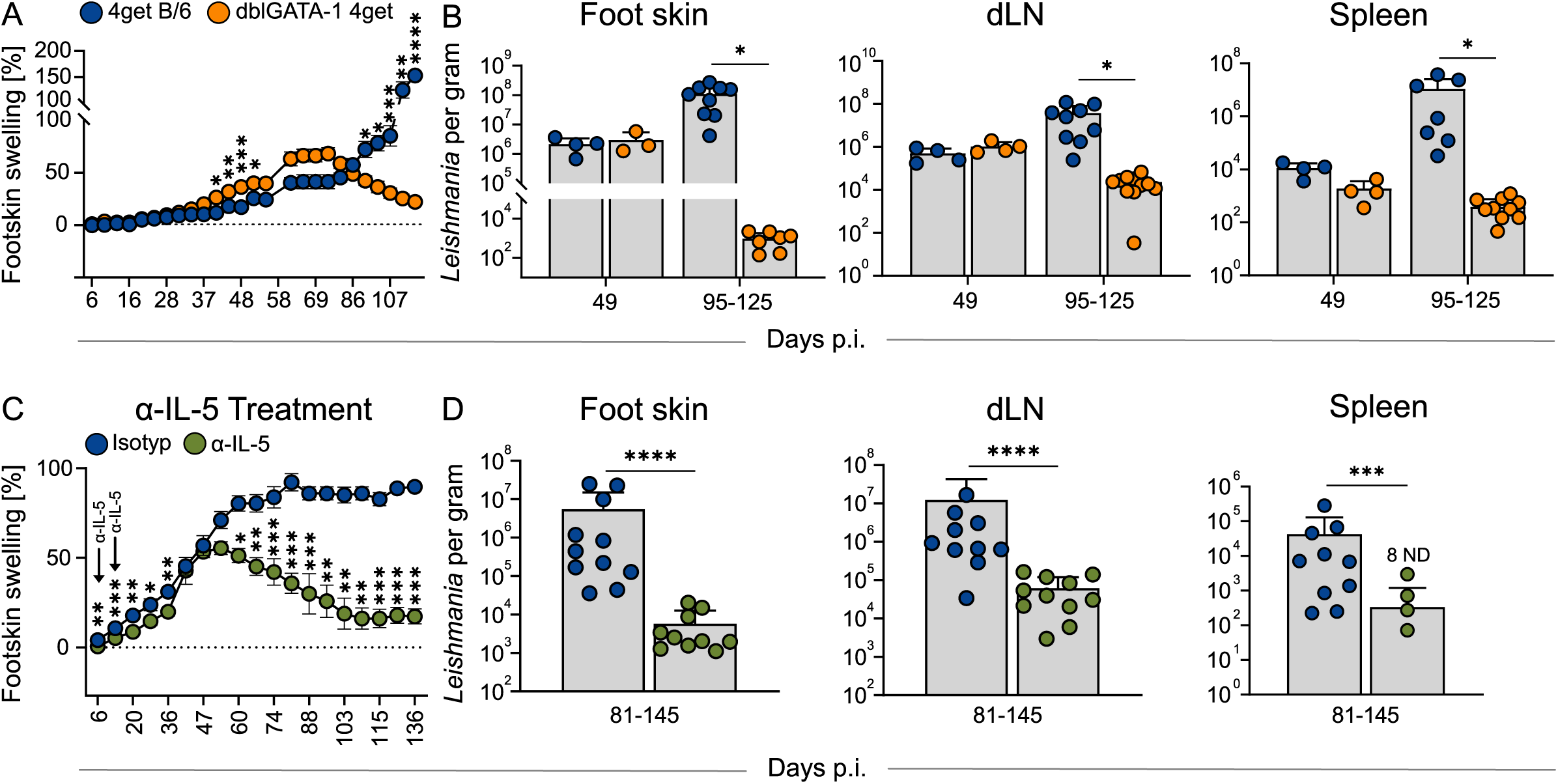
Lack of eosinophils or inhibition of eosinophil expansion prevents chronicity of *L. mexicana* infection **A** Representative clinical course in 4get (WT) and dblGATA-1 4get (KO) mice infected with *L. mexicana* (n > 10 mice per group, 1 of 4 independent experiments). **B** Parasite load determined by two-fold serial dilution of respective organs comparing WT and KO (n = 3-12 mice per group, 1-3 independent experiments). **C** Representative clinical course in C57BL/6 mice infected with *L. mexicana* and treated with 500 µg of either isotype control or anti-IL-5 antibody on days 5 and 12 p.i. (n = 12 mice per group, 1 of 2 independent experiments). **D** Parasite load determined by two-fold serial dilution of the respective organs, comparing isotype control and anti-IL-5 treatment (n = 10-12, 3 independent experiments). *p ≤ 0.05; **p ≤ 0.01; ***p ≤ 0.005; ****p ≤ 0.0001, two-tailed Mann-Whitney U test. Data are mean ± SEM (**A, C**) or mean ± s.d. (**B**). ND: not detectable.

As the dblGATA-1 mutation might cause eosinophil-independent effects during hematopoiesis (Harigae et al., 1998; Hwang et al., 2022; Nei et al., 2013; Pevny et al., 1995; Vyas et al., 1999), we decided to inhibit the expansion and infiltration of eosinophils also by a pharmacological approach and administered a neutralizing anti-IL-5 antibody on days 5 and 12 p.i., which prevented local and systemic eosinophilia (**Figure 1F**). Strikingly, this early IL-5 blockade led to the same self-healing phenotype and reduced parasite burden as seen in the dblGATA-1 mice (**Figure 3C** and **D**). Thus, interruption of the expansion and tissue infiltration of eosinophils in the early phase of infection is sufficient to completely reverse the clinical course of infection.

### *L. mexicana* infection causes a unique inflammatory transcriptomic signature in skin lesion eosinophils

In order to define mechanisms, by which eosinophils fulfil the observed regulatory function in CL, we characterized the transcriptome of circulating and skin-infiltrating eosinophils early after infection (day 14) by performing nanowell-based scRNA-seq of total viable foot skin cells and Percoll-enriched granulocytes of the blood (**Figure S2A**). After *in silico* removal of all non-eosinophil cells, we subjected the remaining sequences of the eosinophil cells to a uniform manifold approximation and projection for dimension reduction (UMAP). This analysis unveiled six distinct eosinophil populations, all of which expressed the canonical eosinophil markers *Il5ra*, *Ccr3*, *Siglecf* and *Prg2* (**Figure 4A, Figure S2B**). Circulating eosinophils predominantly derived from the blood, whereas the remaining clusters originated from the skin (**Figure S2C**). As shown in **Figure 4B, circulating eosinophils** expressed several immediate early response genes, including *Fos*, *Fosb* and *Junb*, which are AP-1 transcription factor family members involved in rapid cellular responses to stimuli (Shaulian and Karin, 2002), as well as *Cd24a*, a recently identified marker for blood-derived eosinophils (Gurtner et al., 2023). **Inflammatory eosinophils I** displayed a similar, though slightly enhanced, expression profile, while concurrently upregulating inflammatory genes, such as *Cd274* (*Pdl1*), a recently identified marker for activated eosinophils (Gurtner et al., 2023), as well as *Il1rn*, *Fcgr3*, *Hif1a*, and *Nfkb1*. Notably, these cells demonstrated a marked upregulation of *Slc2a3*, which is known as the high affinity glucose transporter (Manolescu et al., 2007). **Inflammatory eosinophils II** lost the immediate early response gene signature and downregulated *Slc2a3*, while further intensifying the expression of inflammatory genes. In contrast, **inflammatory eosinophils III** showed reduced expression of both immediate early response and inflammatory genes, while upregulating mitochondrial genes (*mt-Co3*, *mt-Nd1*), suggesting enhanced mitochondrial activity. **Tissue repair eosinophils** exhibited a gene signature similar to inflammatory eosinophils III, while also upregulating genes associated with extracellular matrix remodelling and tissue repair, including *Fbln2*, *Col6a2*, *Adamts2* and *Adamts5* (Loreti and Sacco, 2022; Lu et al., 2011; Pan et al., 1993; Salaiza-Suazo et al., 2024). The sixth cluster termed “**lncRNA eosinophils**” was characterized by significant upregulation of several long non-coding (lnc) RNA genes with yet unknown functions. Together, these data indicate that by day 14 after cutaneous infection, circulating blood eosinophils already acquired a pre-inflammatory gene signature. After infiltration into the infected skin, eosinophils followed a trajectory through distinct transcriptional states and acquired tissue imprinting. This process started with cluster I, the signature of which reflects “priming by recruitment” as well as “early skin-tissue imprinting” with upregulated *Slc2a3.* Inflammatory Eos I then underwent further activation and developed into fully inflammatory eosinophil clusters II and III.

**Figure 4:**
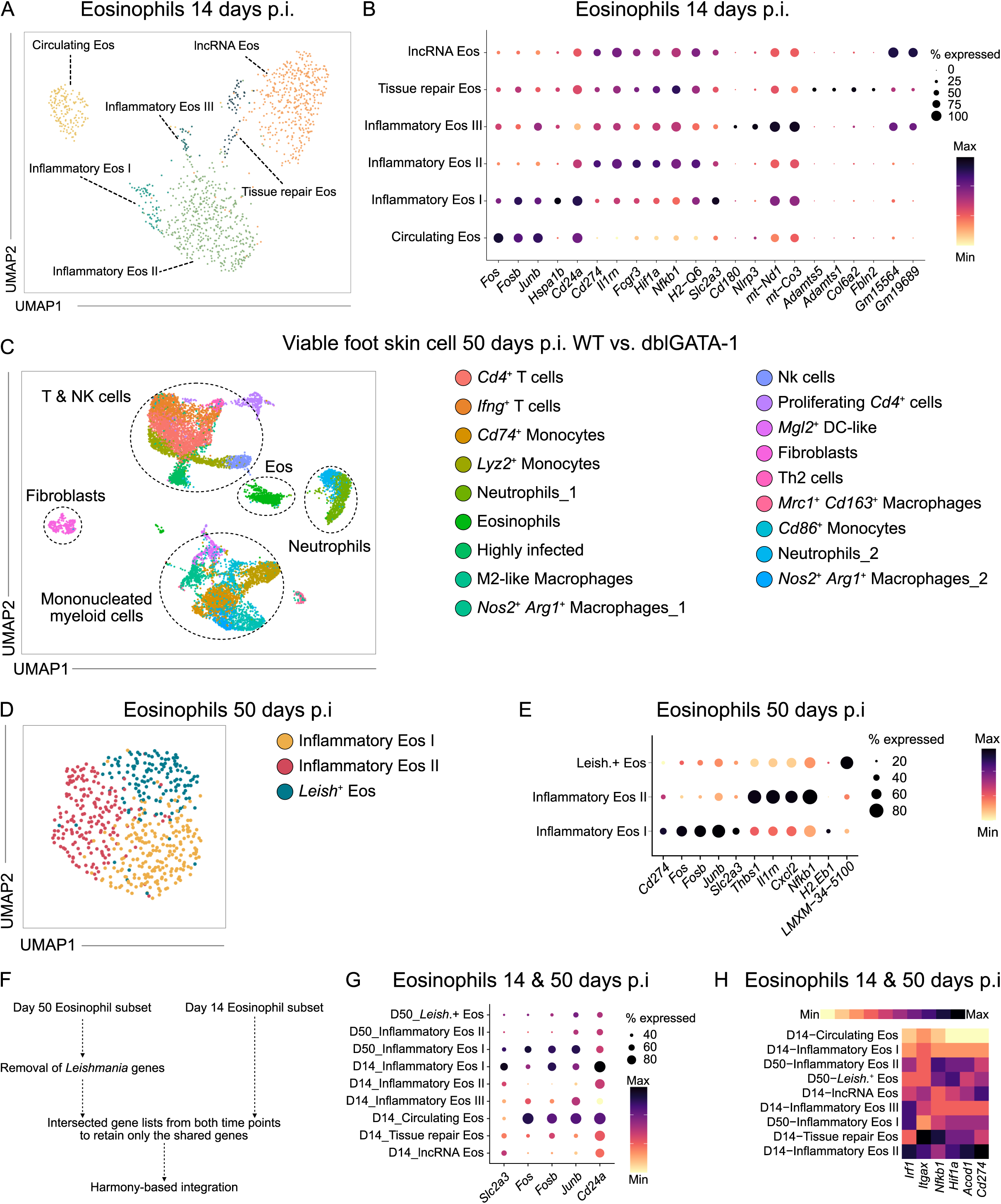
*L. mexicana* infection causes a unique inflammatory transcriptomic signature in skin lesion eosinophils **A** UMAP visualization of skin and blood eosinophils on day 14 p.i. (n = 12 C57BL/6 mice pooled). **B** Expression dot plot showing marker genes that discriminate eosinophil subsets on day 14 p.i. **C** UMAP of total viable skin lesion cells on day 14 p.i. from 4get (WT) and dblGATA-1 4get (KO) mice (n = 7 pooled mice per group). **D** UMAP visualization of skin eosinophils on day 50 p.i. derived from data depicted in (**C**). **E** Expression dot plot of marker genes discriminating the eosinophil subsets on day 50 p.i. **F** Flow chart depicting the computational integration of the eosinophil subsets from day 14 p.i. (**A**) and day 50 p.i. (**D)**. **G** Expression dot plots of selected marker genes, comparing integrated day 14 p.i. and day 50 p.i. eosinophil populations. **H** Average expression heat map of selected activation markers comparing integrated day 14 p.i. and day 50 p.i. eosinophil populations.

Finally, we compared the transcriptomes of blood and skin eosinophils from *L. mexicana* infected mice (day 14 p.i.) with published transcriptomes. Our analysis revealed that the eosinophil signature of the *Clec4a4*^+^ arylhydrocarbon receptor (AHR)-dependent subset of immunomodulatory eosinophils in the small intestine (Wang et al., 2022) was predominantly seen in our circulating eosinophil population, while the signature of *Clec4a4*^-^ eosinophils with a proinflammatory and tissue repair profile (Wang et al., 2022) prevailed in our inflammatory eosinophils II (**Figure S2D**). Integrating the dataset of human liver eosinophils obtained during chronic viral hepatitis C infection (Cui et al., 2024), the UMAP-projection revealed distinct clustering of the eosinophils based on their tissue origin (**Figure S2E, Figure S2F**). The liver eosinophils displayed a transcriptional profile most similar to blood eosinophils, characterized by the expression of the immediate early response genes *Fos*, *Fosb* and *Junb* (**Figure S2F**). From these data, we conclude that *L. mexicana* infection leads to a unique inflammatory signature in skin eosinophils.

To gain further insights into the disease-promoting effects of inflammatory skin eosinophils, we performed a second scRNA-seq analysis on total viable foot skin cells of WT and dblGATA-1 mice at a time-point (day 50 p.i.) (**Figure S3A**), when the clinical disease of WT and dblGATA-1 mice was still comparable (**Figure 3A** and **B**), but the eosinophil number had already dropped in the WT controls (**Figure 1A**). Unbiased clustering analysis revealed a total of 19 distinct clusters, which were comprised of five major cell types: T/NK cells, monocytes/macrophages, fibroblasts, eosinophils and neutrophils (**Figure 4C, Figure S3B-C**). Sub-clustering of the eosinophil population within the total viable skin cells revealed three distinct eosinophil subsets, including the inflammatory eosinophil I and II subset with a similar signature as seen at day 14 p.i. and an additional subset with high expression of *Leishmania* genes, suggesting that eosinophils have internalized the parasite *in vivo* (**Figure 4D** and **E**). The presence of *Leishmania* inside eosinophils was confirmed by sorting of skin eosinophils (SiglecF^+^ Ly6G^-^) 20 days p. i. and immunofluorescence staining of *Leishmania*. Approximately 30% of purified eosinophils contained intracellular parasites (**Figure S3D**, white arrows). We utilized the eosinophil scRNA-seq data from day 50 p.i. to perform a longitudinal data integration with our eosinophil profiles from day 14 p.i. (**Figure 4F**). UMAP-projection after *in silico* removal of all *Leishmania* genes revealed that only the inflammatory eosinophil I and II subsets showed distinct overlap between the two time points (**Figure S3E**). Comparing the characteristic gene signature of the two inflammatory eosinophil I subsets, *Slc2a3* expression was much higher on day 14 p.i. than on day 50 p.i. (**Figure 4G**). Inflammatory eosinophils II displayed a more pronounced inflammatory phenotype on day 14 p.i. compared to day 50 p.i. (**Figure 4H**).

Collectively, these findings strongly suggest that the microenvironment in the *L. mexicana*-infected skin not only leads to the attraction of eosinophils, but also elicits the upregulation of inflammatory genes including the high affinity glucose receptor 3 (*Slc2a3*), which argues for a high glucose demand. In addition, recruited eosinophils represent host cells for *Leishmania*.

### Lack of eosinophils in *L. mexicana*-infected mice induces a shift towards inflammatory macrophages

Considering a possible immunoregulatory role of eosinophils, we next addressed the impact of eosinophil deficiency on various immune cell compartments using scRNA-seq. Comparative analysis of the relative cell abundances in WT versus dblGATA-1 mice at day 50 p.i. illustrated that the absence of eosinophils especially led to an increase in *Nos2*^+^ *Arg1*^+^ macrophages and *Ifng*^+^ Th1 cells as well as to a decrease in *Mrc1*^+^ *Cd163*^+^ macrophages and M2-like macrophages (**Figure 5A** and **B**). Sub-clustering analysis of the monocyte/macrophage population revealed nine distinct clusters (**Figure 5C, Figure S4A**). Direct comparison of the clusters seen in WT and dblGATA-1 mice again demonstrated a nearly complete absence of M2-like macrophages as well as an increase in M1-like macrophages in the eosinophil-deficient animals (**Figure 5D**). Further characterization showed that M2-like macrophages 1 exhibited clear expression of *Retnla*, *Mrc1*^high^ (CD206), *Mgl2* (CD301b) and *Cd163*, whereas M2-like macrophages 2 displayed minimal expression of *Retnla* and *Cd163*, but high *Ccl24* (eotaxin-2) and *Mgl2* (CD301b) and intermediate *Mrc1* (CD206) expression (**Figure 5E**). The prominent expression of *Ccl24* is in line with the rapid infiltration of the skin lesions by eosinophils (**Figure 1 A**), which we found to be a major source of IL-4 in *L. mexicana*-infected mice (**Figure S4B and C**) as previously described for *L. major* infection (Lee et al., 2020; Sasse et al., 2022). Flow cytometry confirmed the significant reduction of SiglecF^-^ Ly6G^-^ CD11b^+^ CD206^+^ CD163^+^ M2-like cells in *L. mexicana*-infected skin of dblGATA-1 mice as compared to WT mice (**Figure 5F**). In addition, the absence of eosinophils was associated with an expansion of M1-like macrophages characterized by the expression of *Nos2* (**Figure 5G**). Although most of the M1-like macrophages appeared to co-express *Nos2* and *Arg1* in WT and dblGATA-1 mice (**Figure 5G**), RT-qPCR analysis of total skin lysates yielded an increased expression of *Nos2* at day 7 to 48 p.i. in dblGATA-1 mice as compared to WT mice, whereas the overall mRNA expression of *Arg1* differed only weakly between both mouse strains (**Figure 5H**). Together, these findings indicate that depletion of eosinophils induces a shift towards a more inflammatory macrophage phenotype during *L. mexicana* infection.

**Figure 5:**
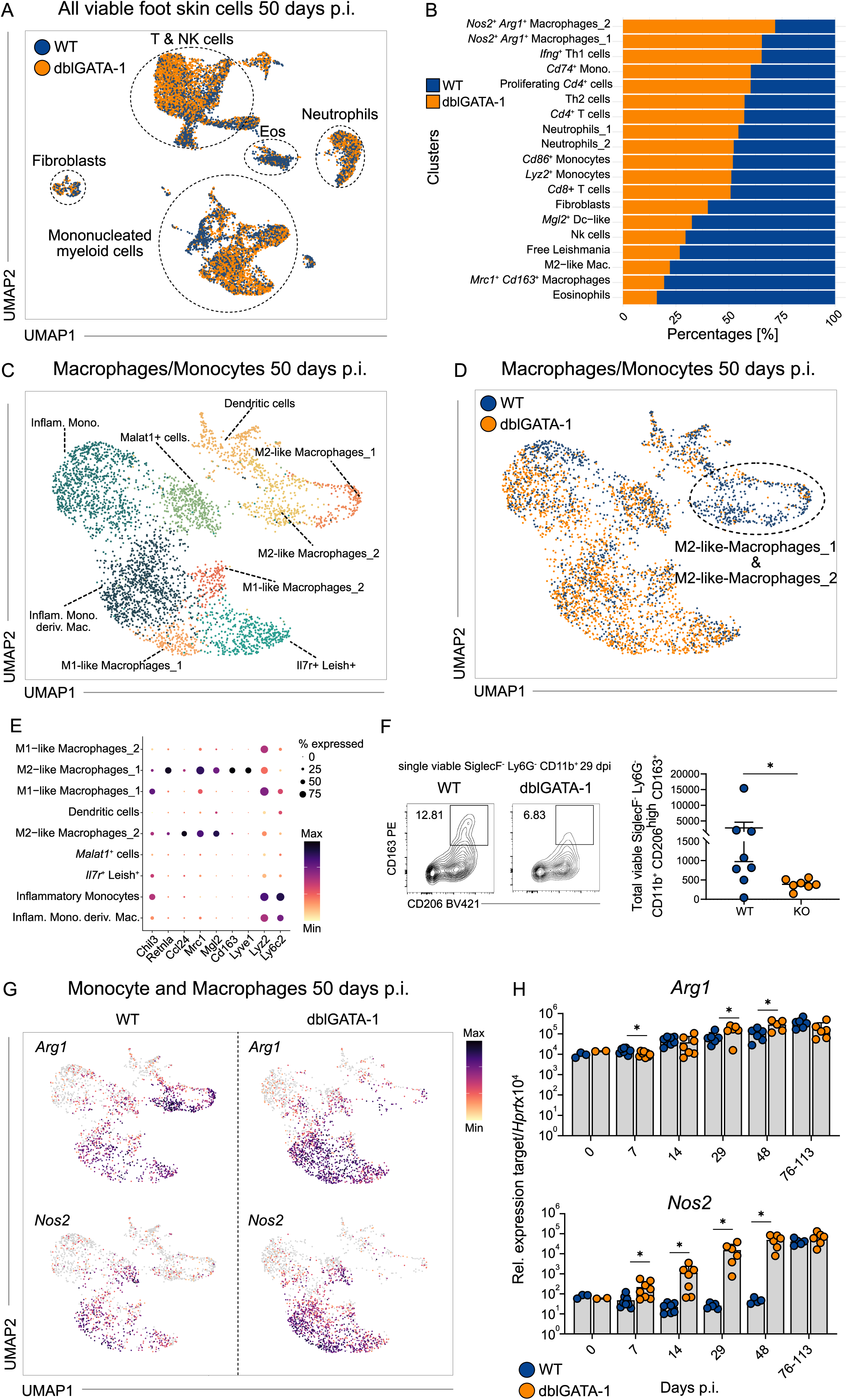
Lack of eosinophils in *L. mexicana*-infected mice induces a shift towards inflammatory macrophages **A** UMAP of total viable skin lesion cells at day 50 p.i. from 4get (WT) and dblGATA-1 4get (KO) mice (n = 7 pooled mice per group). **B** Stacked bar plot comparing relative abundance of WT and KO in each macrophage/monocyte cluster in (**A**). **C**/**D** UMAP of the monocyte and macrophage clusters derived from (**A**). **E** Expression dot plot of marker genes identifying M2-like macrophages_1 and M2-like macrophages_2. **F** Left, representative flow cytometric analysis of the M2-like macrophages_1 in skin lesions at day 29 p.i.. Right, quantification of the respective WT and KO populations (n = 7-8 mice, 2 independent experiments). **G** Feature plots displaying the expression of selected markers overlaid on the UMAP on day 50 p.i.. Expression intensity is represented by a colour gradient, with grey indicating no detectable expression. **H** Quantification of arginase (*Arg1*) and nitric oxide synthase 2 (*Nos2*) mRNA expression by qRT-PCR in *L. mexicana*-induced skin lesions (n = 2-8 mice per time point, 2 independent experiments). *p ≤ 0.05, two-tailed Mann-Whitney U test. Data are mean ± SEM **(F)** or mean ± s.d. **(H)**.

### Eosinophil depletion in *L. mexicana* infection promotes expansion and activation of Th1 cells

As indicated in **Figure 5B**, expansion of *Nos2*^+^ macrophages was paralleled by an increase in *Ifng^+^* Th1 cells in infected dblGATA-1 mice. We therefore conducted a detailed comparison of the T/NK cell clusters in WT and dblGATA-1 mice. Unbiased sub-clustering of these cells revealed seven distinct cell types (**Figure 6A, Figure S4D**). Six clusters were identified as NK cells, CD8^+^ T cells, central memory T cells (TCM), γδ T cells, Th2 cells and as *Ifng*^+^ Th1 cells. In addition, we found a T cell cluster characterized by the presence of *Leishmania* transcripts (Leish^+^), which presumably results from the incorporation of parasite mRNA through interactions with (debris or extracellular vesicles of) infected cells, although endocytosis of *Leishmania* amastigotes by (activated) T cells cannot formally be excluded (Wu et al., 2009). Besides the substantial increase in *Ifng*^+^ Th1 cells, there was also a minor expansion of Th2 cells in the eosinophil-deficient mice (**Figure 6B**). Flow cytometry of skin lesions confirmed the increased infiltration of activated CD4^+^CD44^+^ T cells in the *L. mexicana*-infected dblGATA-1 mice (**Figure 6C**). The enhanced expression of IFNγ by Th1 cells in the absence of eosinophils was not only seen in the scRNA-seq (**Figure 6D**), but was also corroborated by RT-qPCR analysis (**Figure 6E**) and by intracellular IFNγ staining of CD11b^-^ CD3^+^ CD4^+^ cells (**Figure 6F**) performed with total skin cell lysates or cell suspensions, respectively. While we also observed a slight increase in *Il4* expression within the Th2 cluster (**Figure S4E**) and in whole skin lysate (**Figure 6E**), the enhancement of the Th1 response was substantially more pronounced. The improved T cell response in the dblGATA-1 mice was also reflected at the metabolic level, with a notable upregulation of a glycolytic gene signature especially in the Th1 cluster (**Figure 6G**). From these data we conclude that depletion of eosinophils strengthens the Th1 response after *L. mexicana* infection.

**Figure 6:**
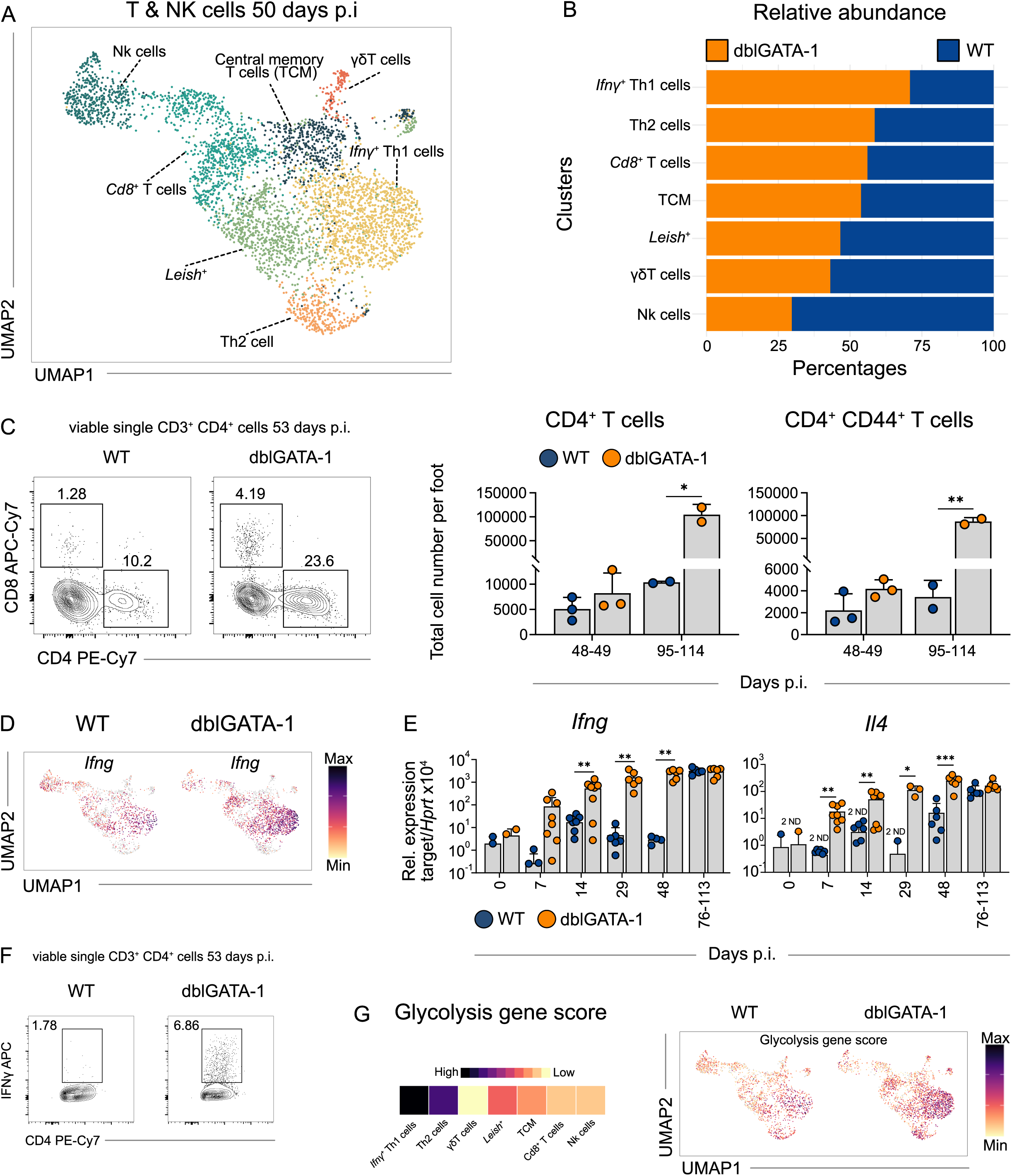
Eosinophil depletion in *L. mexicana* infection promotes expansion and activation of Th1 cells **A** UMAP of the T- and NK cell clusters derived from Figure 4C (total viable cells from foot skin lesion at day 50 p.i.). **B** Stacked bar plot comparing relative abundance of 4get (WT) and dblGATA-1 4get (KO) in each T cell and NK cell cluster in (**A**). **C** Left, representative flow cytometric analysis of CD4^+^ T cells in skin lesions at day 53 p.i.. Right, quantification of total CD4^+^ and CD4^+^ CD44^high^ T cell numbers using flow cytometry (n = 2-3, data points were derived from samples pooled from two mice, 2 independent experiments). **D** Feature plots displaying the expression of *Ifng* overlaid on the UMAP of T and NK cell subset. Expression intensity is represented by a colour gradient, with grey indicating no detectable expression. **E** Quantification of *Ifng* and *Il4* mRNA expression by qRT-PCR in *L. mexicana*-induced skin lesions (n = 2-8 mice per time point, 2 independent experiments). **F** Representative flow cytometric analysis of IFNγ-producing T cells in skin lesions at day 53 p.i. (n = 6, 2 independent experiments). **G** Glycolysis gene score extracted from Gene Ontology ID: 0006006 visualized as a heatmap (left, not discriminating between WT and KO) and a feature plot (right, comparing WT vs KO). **(C, D)** *p ≤ 0.05; **p ≤ 0.01; ***p ≤ 0.005, two-tailed Mann-Whitney U test. Data are mean ± s.d.

### Eosinophils derived from *L. mexicana*-infected skin directly repress the Th1 response

Activated effector T cells are strongly glycolytic (**Figure 6G**) and have a high need for glucose uptake and metabolism for full function (Buck et al., 2015; Chang et al., 2013; Ma et al., 2019). We therefore analyzed the expression of glucose transporters using our scRNA-seq data obtained from WT mice at day 50 p.i.. Intriguingly, besides skin-imprinted inflammatory eosinophils, *Ifng*^+^ Th1, Th2 and proliferating *Cd4*^+^ cells were the only cell clusters that expressed the high affinity glucose transporter *Slc2a3* (**Figure 7A**), whereas the more abundant glucose transporter 1 (*Slc2a1*) demonstrated a much broader distribution across various cell types (**Figure S5A**). Of note, eosinophils showed the highest expression of *Slc2a3* (**Figure 7A**). Based on these observations, we hypothesized that the early influx of eosinophils might suppress T cell responses in the skin by restricting their glucose uptake and that Th1 cells are particularly affected, because they showed the highest glycolytic profile (**Figure 6G**). To test this hypothesis, we isolated all viable skin cells at day 20 p.i. and incubated them with the fluorescently labelled glucose analogue 2-NBDG (**Figure 7B**). Flow cytometry revealed that eosinophils (CD11b^+^ SiglecF^+^ Ly6G^-^) took up 2-NBDG at a much higher percentage than monocytes/macrophages (CD11b^+^ SiglecF^-^ Ly6G^-^) or CD3^+^ CD4^+^ T cells (**Figure 7C** and **D**). Next, we investigated whether eosinophils have a direct effect on the activation of Th1 cells. When eosinophils sorted from skin lesions at day 20 p.i. (SSC^high^ SiglecF^+^ Ly6G^-^) were co-cultivated in a 1:1 ratio with *in vitro* generated Th1 cells overnight (**Figure 7E**), the mean fluorescence intensity (MFI) of CD44^+^ and the production of IFNγ were significantly reduced compared to control cultures without eosinophils (**Figure 7F, Figure S5B**). *In vivo*, this effect is likely to be even more pronounced, because skin lesions contained 12.4 (α 2.8)-fold more eosinophils than CD4^+^ T cells at day 20 p.i. (mean +/- SD of 9 lesions from two experiments; **Figure S5C**). Together, these findings strongly indicate that inflammatory eosinophils expressing high levels of *Slc2a3* deprive activated Th1 cells of glucose in the *L. mexicana*-infected skin and thereby inhibit an efficient and protective Th1 response during the early phase of infection.

**Figure 7:**
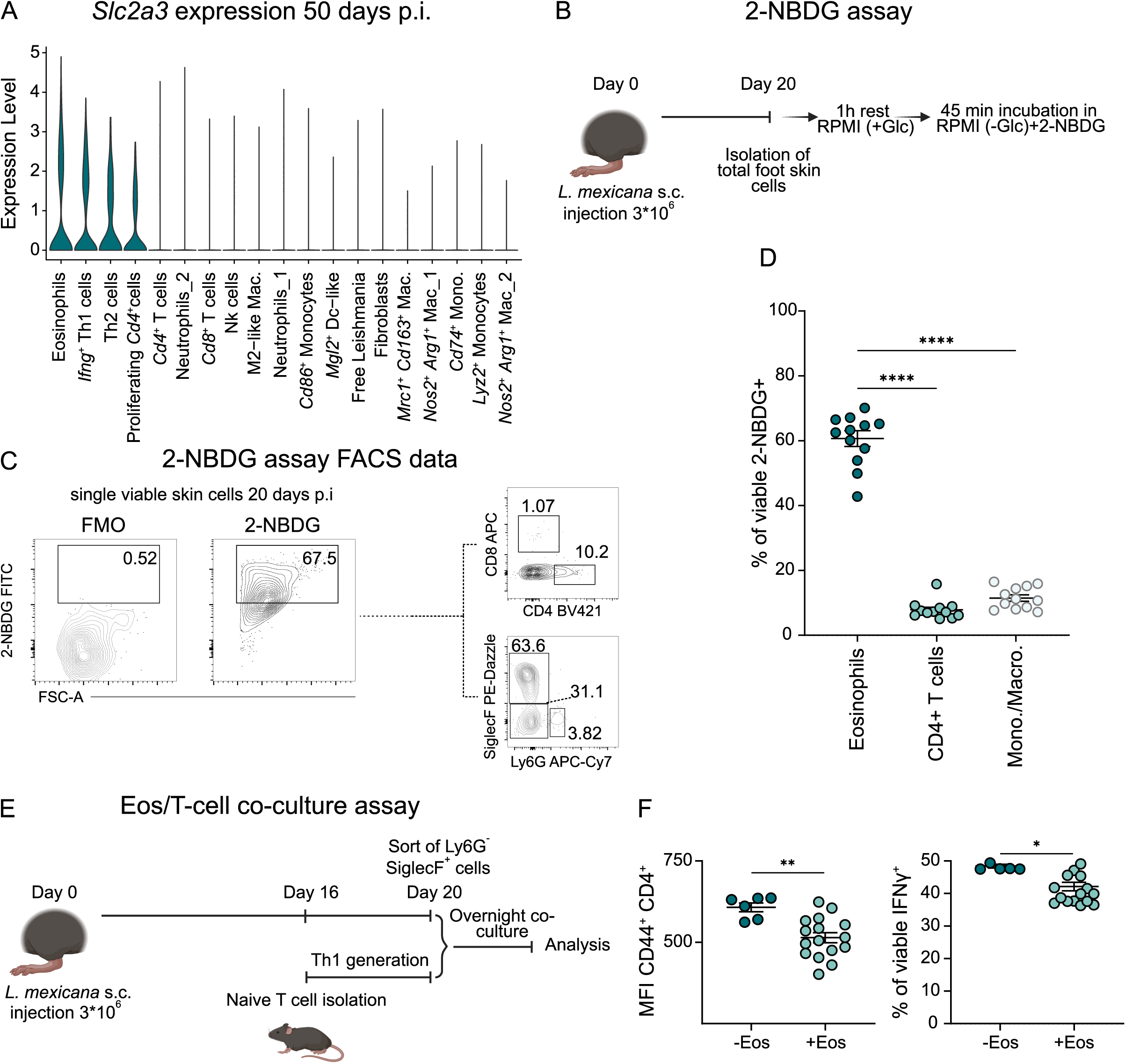
Eosinophils derived from *L. mexicana*-infected skin directly repress the Th1 response **A** Violin plot depicting the distribution of *Slc2a3* (solute carrier family 2 member 3, glucose transporter 3) expression across all cluster of total viable skin lesion cells at day 50 p.i.. **B** Experimental workflow of the 2-NBDG assay (Glc, glucose). **C** Representative flow cytometric analysis of 2-NBDG uptake by *ex vivo* cultured cells from day 20 p.i.. **D** Quantification of 2-NBDG^+^ eosinophils (CD11b^+^ SiglecF^+^ Ly6G^-^), monocytes/macrophages (CD11b^+^ SiglecF^-^ Ly6G^-^) and CD4^+^ T cells (CD3^+^ CD4^+^). **E** Experimental workflow of the skin eosinophil/T-cell co-culture assay. **F** Flow cytometric quantification of the skin eosinophil/T-cell co-culture assay using the median fluorescence intensity (MFI) of CD44 and the intracellular production of IFNγ of CD4^+^ T cells. Co-cultures were stimulated with phorbol-12-myristate-13-acetate (PMA) and ionomycin for 4h (n = 4-15, 3 independent experiments). **(D, F)** *p ≤ 0.05; **p ≤ 0.01; ****p ≤ 0.0001, two-tailed Mann-Whitney U test. Data are mean ± SEM.

## DISCUSSION

In the past, the function of eosinophils during homeostasis, infections, malignant and allergic diseases has been mostly related to their ability to express antimicrobial and cytotoxic pathways or to release immunoregulatory cytokines and chemokines (Arnold et al., 2018; Jacobsen et al., 2021; Kanda et al., 2020; Wen and Rothenberg, 2016). In the present study, we demonstrate that eosinophils do not act as critical antimicrobial effector cells, but instead have a fundamental negative impact on the course and outcome of a protozoan skin infection. We provide evidence that the disease-promoting effect of eosinophils is related to their superiority in the uptake of glucose and the consecutive inhibition of Th1 function and not just to their production of IL-4. We also present a comprehensive transcriptomic analysis of circulating as well as skin-infiltrating eosinophils in this model of chronic CL. Together, our findings illustrate the prominent influence of the tissue immunomicrotope (Bogdan et al., 2024) on the transcriptome and function of eosinophils and highlight a novel immunoregulatory activity of these innate immune cells.

### Eosinophils and *Leishmania*

Earlier studies, which investigated the role of eosinophils in the pathogenesis of CL, were either limited to the mere detection of eosinophils infiltrating the skin lesions (Beil et al., 1992; Grimaldi et al., 1984; McElrath et al., 1987; Pompeu et al., 1991) or described an antileishmanial effector function of eosinophils (Almeida et al., 2024; da Silva Marques et al., 2021; Oliveira et al., 1998; Pimenta et al., 1987; Salaiza-Suazo et al., 2024; Watanabe et al., 2004). Recent research in CL has focused on immunomodulatory effects of eosinophils. In a mouse model of progressive CL using the *L. major* Seidman strain, Lee *et al*. unravelled a regulatory circuit, in which TRM released CCL24 that led to the recruitment of eosinophils, as well as TLSP that activated ILC2 for the production of IL-5 and thereby further promoted the expansion of eosinophils. Eosinophils, in turn, formed a major source of IL-4, which converted the dermal TRMs into a M2-like macrophage population serving as a replicative niche for the parasite (Lee et al., 2020; Lee et al., 2023). In *L. mexicana*-infected mice, we similarly observed an ILC2- and IL-5-dependent eosinophilia, an expression of CCL24 by M2-like macrophages in the skin lesion and a prominent production of IL-4 by eosinophils (**Figures 5E, S4B, S4C**). However, there are also remarkable differences between the two models. First, a disruption of the above-mentioned regulatory circuit by deletion of eosinophilic IL-4, of IL-5^+^ ILC2 or of TLSP in CCL24-expressing TRM only caused an amelioration of the course of *L. major* Seidman infection (Lee et al., 2020; Lee et al., 2023). In contrast, genetic or pharmacological depletion of eosinophils in *L. mexicana*-infected mice completely reverted the disease outcome and allowed for clinical healing (**Figures 3A** and **3C**). Second, in eosinophil-depleted *L. mexicana*-infected mice, we still observed an upregulation of IL-4 in the skin lesion, both in our scRNA-seq data (**Figure S4B and S4C**) and in our RT-qPCR analyses (**Figure 6E**). Our findings strongly suggest that infiltrating, inflammatory eosinophils are the drivers of chronic CL and that the disease-promoting effect of eosinophils is not primarily due to the production of IL-4. While eosinophil-as well as T-cell-derived IL-4 upregulate ARG1 expression (**Figure 5G** and **5H**), which antagonizes the activity of NOS2 in macrophages and is critical for generating a replicative host cell niche for *Leishmania* parasites (Bogdan, 2020; Bogdan et al., 2024; Schleicher et al., 2016), the detrimental effect of eosinophils in *L. mexicana*-infected mice appears to result also from their metabolic signature (**Figure 7A**) as discussed below.

The accumulation of eosinophils at the skin site following infection with *L. mexicana* has not only been observed in mice, but also in humans (Grimaldi et al., 1984; McElrath et al., 1987; Salaiza-Suazo et al., 2024). Salaiza-Suazo *et al*. compared eosinophil responses in patients with localized (LCL) and diffuse cutaneous leishmaniasis (DCL). Whereas all DCL patients exhibited blood eosinophilia and elevated eosinophil counts in non-ulcerated nodules, only LCL patients with a prolonged course of disease developed blood eosinophilia and cutaneous ulcers (Salaiza-Suazo et al., 2024). Thus, similar to our mouse model, the severity of human CL appears to correlate with the extent of eosinophil activation. Furthermore, it is tempting to speculate that the massively impaired IFNγ response in DCL patients (Bomfim et al., 1996) might at least partially result from the prominent eosinophilia in these patients.

### Eosinophils, metabolism and regulation of T cell activity

A number of previous studies have documented unexpected regulatory effects of eosinophils on T lymphocytes. In cancer models, eosinophils were found to enhance the tumor-infiltrating (Carretero et al., 2015) or anti-tumor activity of CD8^+^ T cells (Blomberg et al., 2023). In mice infected with *Listeria monocytogenes*, eosinophil-derived IL-4 supported the survival and memory cell formation in the CD8^+^ T cell compartment (Zhou et al., 2024), whereas in mice colonized with the gastric pathogen *Helicobacter pylori* IFNγ-activated eosinophils expressed PD-L1 and thereby restrained Th1 responses to maintain gastrointestinal homeostasis (Arnold et al., 2018). In the present study, we provide evidence for a novel mechanism of suppression of Th1 effector cell activity by eosinophils. Both eosinophils and T cells have a high demand of glucose and are dependent on glucose metabolism (Chang et al., 2013; Porter et al., 2018). Our scRNA-seq analyses revealed a prominent glycolytic signature of the IFNγ^+^ Th1 cells in *L. mexicana*-infected mice (**Figure 6G**). Furthermore, skin lesion-derived eosinophils expressed high amounts of the glucose transporter *Slc2a3*, which exceeded the levels seen in IFNγ^+^ Th1 cells (**Figure 7A**), and showed a much higher frequency in the uptake of glucose compared to T cells and monocytes/macrophages (**Figure 7D**). We therefore propose that eosinophils deprive Th1 cells of glucose and thereby impede their IFNγ production and the resolution of *L. mexicana* infections. This notion is supported by previous data showing that in CD4^+^ T lymphocytes aerobic glycolysis was required for optimal IFNγ production, whereas in the absence of glucose the glycolytic enzyme glycerinaldehyde-3-phosphate-dehydrogenase (GAPDH) functioned as a repressor of IFNγ mRNA translation (Chang et al., 2013). In an unrelated mouse model of tumor metastasis, ILC2s also drove eosinophilia, which then impaired the IFNγ production and cytotoxic activity of NK cells, presumably via modulation of the glucose metabolism (Schuijs et al., 2020). Therefore, the competition between eosinophils and other immune cells for glucose is likely to represent a general immunoregulatory mechanism of eosinophils.

### Eosinophils and functional subsets based on transcriptomic profiling

Our transcriptomic profiling revealed a previously uncharacterized inflammatory transcriptional trajectory in skin-infiltrating eosinophils. The eosinophil population in the peripheral blood expanded following infection and already showed a pre-activated state. After entering the skin, eosinophils adapted to the local tissue environment by up-regulating the high-affinity glucose transporter *Slc2a3* and by gradually increasing their inflammatory profile. A careful comparison of our eosinophil dataset with previous bulk RNA-seq data (Wang et al., 2022) and the limited number of published scRNA-seq profiles of eosinophils (Chhiba and Kuang, 2024; Cui et al., 2024; Gurtner et al., 2023) revealed that eosinophils accumulating in the skin in response to *L. mexicana* had a unique transcriptomic phenotype, although they showed certain similarities to inflammatory eosinophils of the gastrointestinal tract. Gurtner *et al*. (Gurtner et al., 2023) identified two distinct subsets of gastrointestinal eosinophils in *Il5*-transgenic mice: active eosinophils (A-Eos) and basal eosinophils (B-Eos). A-Eos developed from B-Eos in response to bacterial cues, IL-33 and IFNγ signaling and displayed antimicrobial and immunoregulatory properties. Our skin-infiltrating inflammatory eosinophil II subset shared a distinct transcriptional overlap with A-Eos, with upregulation of genes such as *Cd274*, *Il1rn*, *Hif1a* and *Nfkb1*. However, our remaining clusters did not align closely with the subsets reported by Gurtner *et al*. (Gurtner et al., 2023). The *Clec4a4*^+^ intestinal eosinophil cluster described by Wang *et al*. (Wang et al., 2022) was predominantly seen in our circulating eosinophil population, while the signature of *Clec4a4*^-^ eosinophils prevailed in the inflammatory eosinophils II. The other eosinophil subsets from the skin were again not represented in the comparison. Additionally, Dolitzky *et al*. (Dolitzky et al., 2021) proposed that eosinophils polarize into distinct “type 1” and “type 2” states under different cytokine conditions. Given that *L. mexicana* elicits a mixed Th1 and Th2 immune response (Castellano et al., 2009; Scott and Novais, 2016), eosinophils may acquire both activation states, explaining the similarities and differences in our dataset compared to previous reports. Our scRNA-seq data extend the concept that BM, blood and other tissue eosinophils differ by distinct transcriptional profiles and that eosinophils adapt to their specific niche of residence by “transcriptional maturation” that depends on microenvironmental cues. While this concept has mainly been based on observations made in the gut under homeostatic conditions (Arnold and Munitz, 2024), the present study now provides first data from the skin following infection with *L. mexicana*.

In summary, our analyses have not only unravelled the mechanism underlying the chronic course of *L. mexicana* infection, but also defined a new regulatory effect of eosinophils on Th1 cells. We believe that these data will help in developing novel intervention strategies for severe CL manifestations in humans and enhance our understanding of various cutaneous eosinophil-associated diseases.

## ACKNOWLEDGEMENTS

This research was supported by the Deutsche Forschungsgemeinschaft (DFG) (Research Training Group 2740 “ImmunoMicroTope”, projects A2 to JJ, A4 to SW, A6 to US, A7 to DV and B2 to CB and DL; priority program SPP1937 “Innate lymphoid cells”, grants SCHL 615/1-1 and 1-2 to US and grants BO 996/5-1 and 5-2 to CB; grants BO996/7-1 and SCHL615/3-1 to CB and US), the Interdisciplinary Center for Clinical of Research (IZKF) of the Universitätsklinikum Erlangen (project A87 to US) and the Collaborative Research Center CRC1181 (DFG, project C04 to US and CB). We are grateful to Sigrid Roberts (School of Pharmacy, Pacific University Oregon, U.S.A.) for providing the *L. mexicana* strain. We also thank Thomas Winkler and Sebastian Zundler (Erlangen, Germany) for supplying Rag1^-/-^ mice, E.S. Sánchez Quant (BD Biosciences, Heidelberg, Germany) for guidance on the handling of the BD Rhapsody system, Dr. Arif Ekici as well as Uwe Appelt and Markus Mroz from the University Hospital Erlangen core units “Next Generation Sequencing” and “Cell Sorting and Immunomonitoring”, respectively, for their technical support, and the personnel of the Franz-Penzoldt Preclincal Experimental Animal Center for their professional mouse care. This study is part of the doctoral thesis of David Barinberg to obtain a Dr. rer. nat. degree.

## Author contributions statement

DB, CB and US designed the experiments. DB, HS, TG, BR and DR performed and analysed the experiments. DB carried out the bioinformatic analyses. DL, AUA and MC gave guidance on the bioinformatic analyses. DV, JJ, and SW provided crucial mouse lines and/or offered valuable advice. DB, US and CB wrote the manuscript. All authors read, edited and approved the article.

## Competing interest statement

The authors declare no competing interests.

---

**Figure S1:**
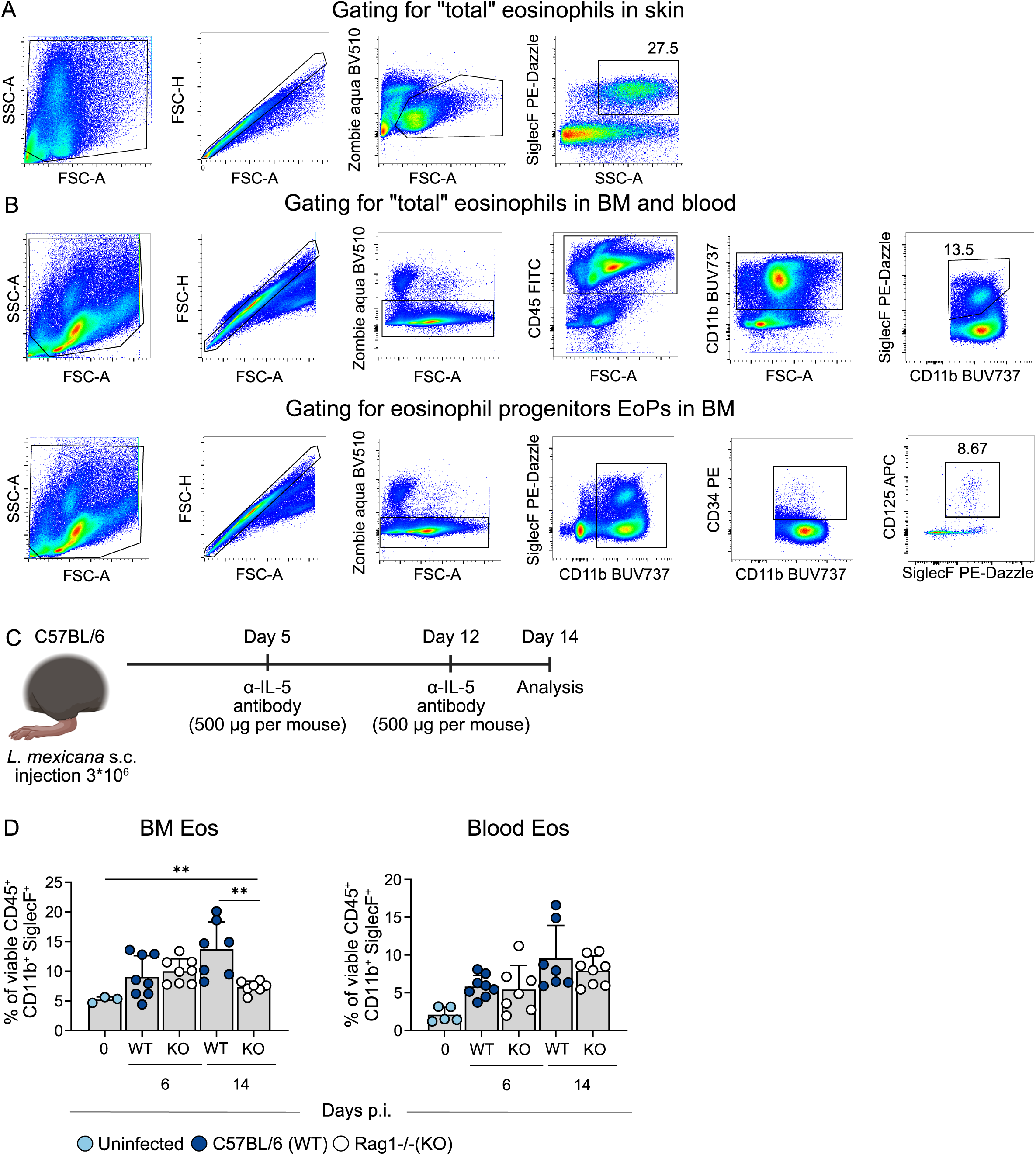
Representative gating strategy for skin (**A**) and BM eosinophil (**B**), as well as eosinophil progenitors (**C**, EoP) at day 14 p.i.. **C** C57BL/6 mice were intraperitoneally treated with 500 µg of either isotype control or anti-IL-5 antibody on days 5 and 12 p.i.. Flow cytometric analysis was performed on day 14 p.i on bone marrow, blood and skin lesions cells. **D** Flow cytometric analysis of cells from bone marrow, blood and skin lesions of C57BL/6 (WT) and Rag1^-/-^ (KO) mice infected with *L. mexicana* (n = 3-8 mice per group, 1-2 independent experiments). **p ≤ 0.01, two-tailed Mann-Whitney U test. Data are mean ± s.d.

**Figure S2:**
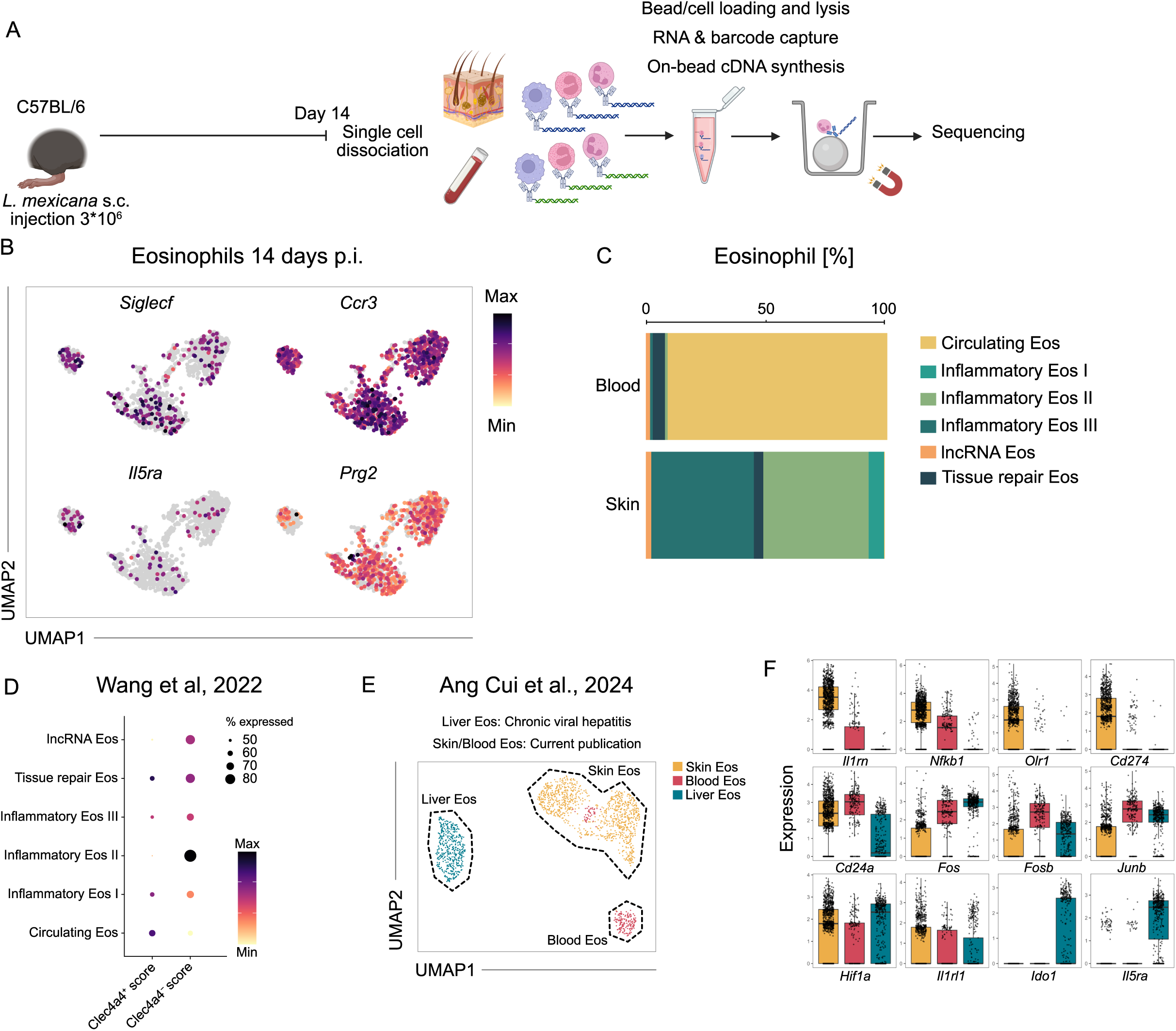
**A** Experimental workflow of nanowell-based scRNA Seq of total viable foot skin cells and Percoll-enriched granulocytes of the blood on day 14 p.i.. **B** Feature plots of canonical eosinophil markers. Expression intensity is represented by a colour gradient, with grey indicating no detectable expression. **C** Relative abundance of eosinophil subsets across blood and skin during infection, as assessed by scRNA-seq. **D** Expression dot plot depicting gene signatures of Clec4a4^+^ and Clec4a4^-^ eosinophil subsets based on published bulk RNA sequencing data from intestinal eosinophils of naïve mice (Wang et al., 2022). **E** UMAP visualization of the computational integration of the eosinophil subsets from day 14 after *L. mexicana* infection (**B**) with liver eosinophils of hepatitis C patients (Cui et al., 2024). **F** Boxplot with jitters representation of differentially expressed genes among eosinophils from the computational integration depicted in **E**.

**Figure S3:**
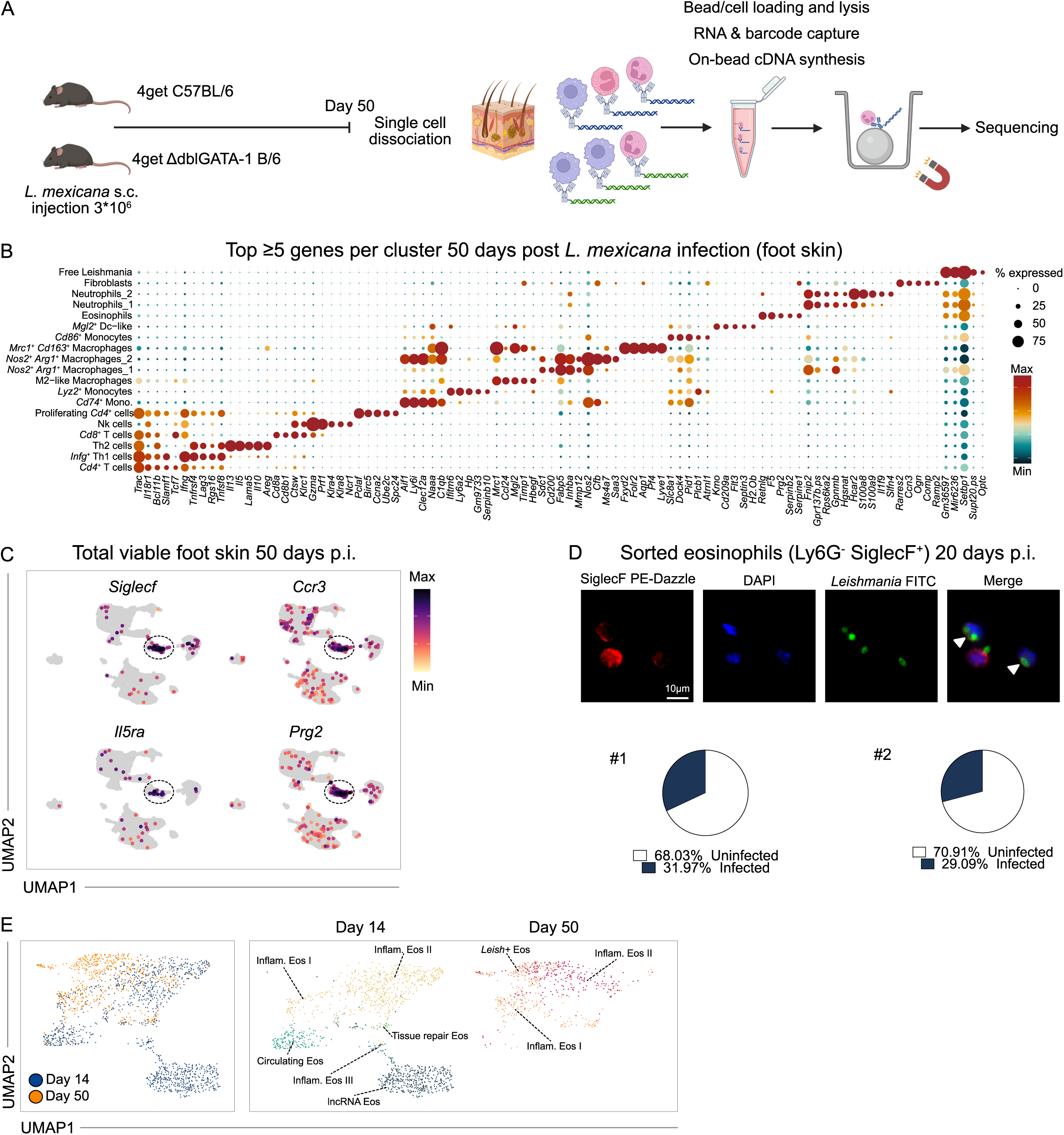
**A** Experimental workflow of nanowell-based scRNA Seq of total viable foot skin cells from 4get C57BL/6 and 4get ΔdblGATA-1 B/6 mice on day 50 p.i.. **B** Expression dot plot depicting the top ≥5 differentially expressed genes for each cluster on day 50 p.i. (total viable foot skin cells). **C** Feature plots of canonical eosinophil markers. Expression intensity is represented by a colour gradient, with grey indicating no detectable expression. The encircled areas represent the eosinophils. **D** Top: representative images of sorted eosinophils (Ly6G^-^ SiglecF^+^; 20 days p.i) stained with anti-SiglecF, anti-*Leishmania* and DAPI. Scale bar, 10 µm. Bottom: quantification of the relative abundance of infected eosinophils (n = 2 independent experiments, 5 mice pooled each, 200 cells counted per quantification). **E** UMAP visualization illustrates the *in silico* integration of eosinophil transcriptomes from day 14 p.i. (see Figure 4A) and day 50 p.i. (see Figure 4D) (see also flow chart Figure 4F).

**Figure S4:**
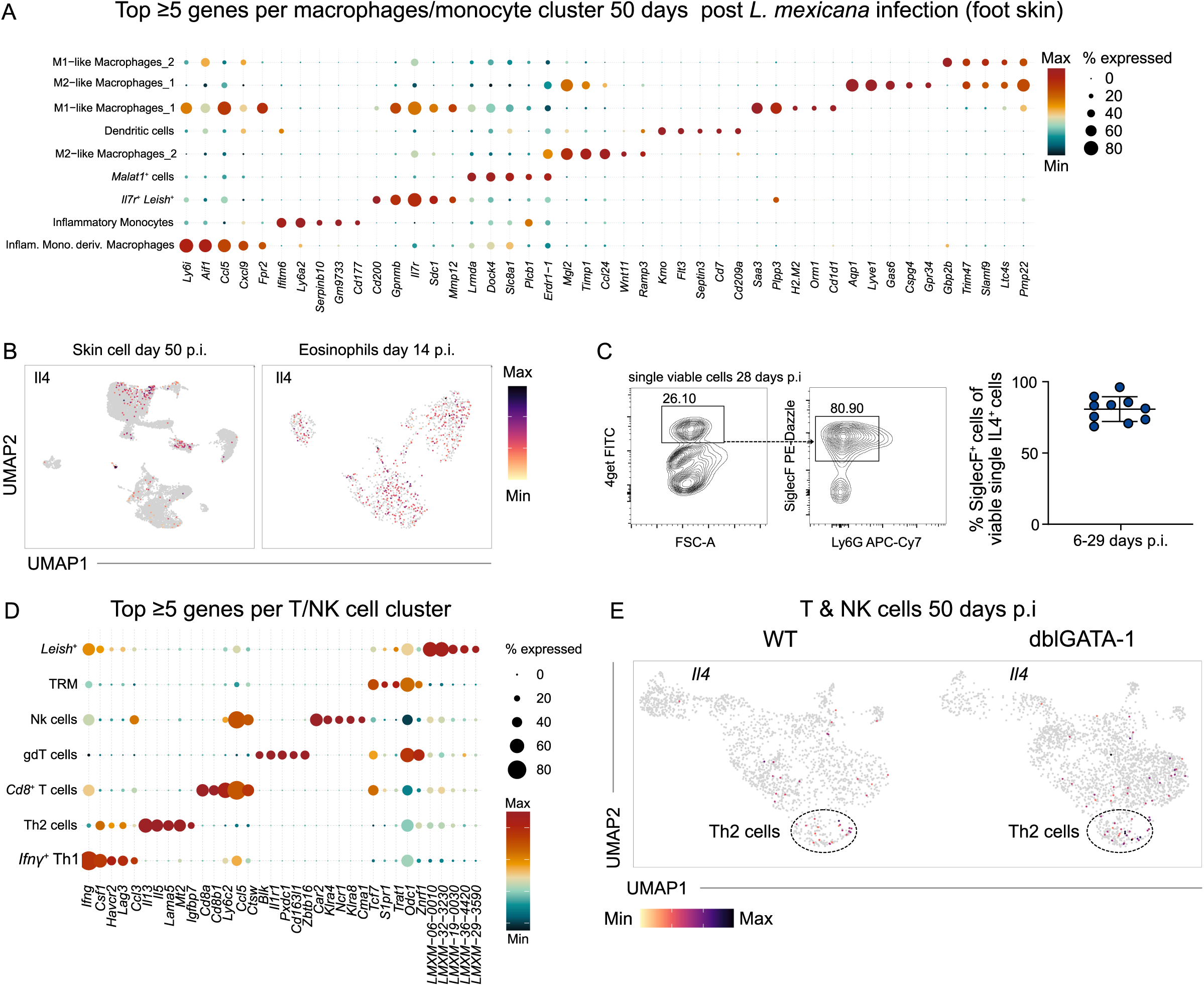
**A** Expression dot plot depicting the top ≥5 differentially expressed genes for each cluster on day 50 p.i. (total viable foot skin). **B** Feature plots displaying the *Il4* expression overlaid on the UMAP on day 50 p.i. (left) and day 14 p.i. (right). Expression intensity is represented by a colour gradient, with grey indicating no detectable expression. **C** Left, representative flow cytometric analysis of 4get^+^ (IL-4) SiglecF^+^ cells at day 28 p.i. Right, relative abundance of SiglecF^+^ cells within the 4get^+^ (IL-4^+^) population of total viable foot skin cells at days 6-29 p.i. (n = 10, 4 independent experiments, mean ± s.d.). **D** Expression dot plot depicting the top ≥5 differentially expressed genes for each T- and NK-cell cluster on day 50 p.i. (total viable foot skin cells). **E** Feature plots displaying the expression of *Il4* overlaid on the UMAP of the T- and NK-cell cluster. Expression intensity is represented by a colour gradient, with grey indicating no detectable expression.

**Figure S5:**
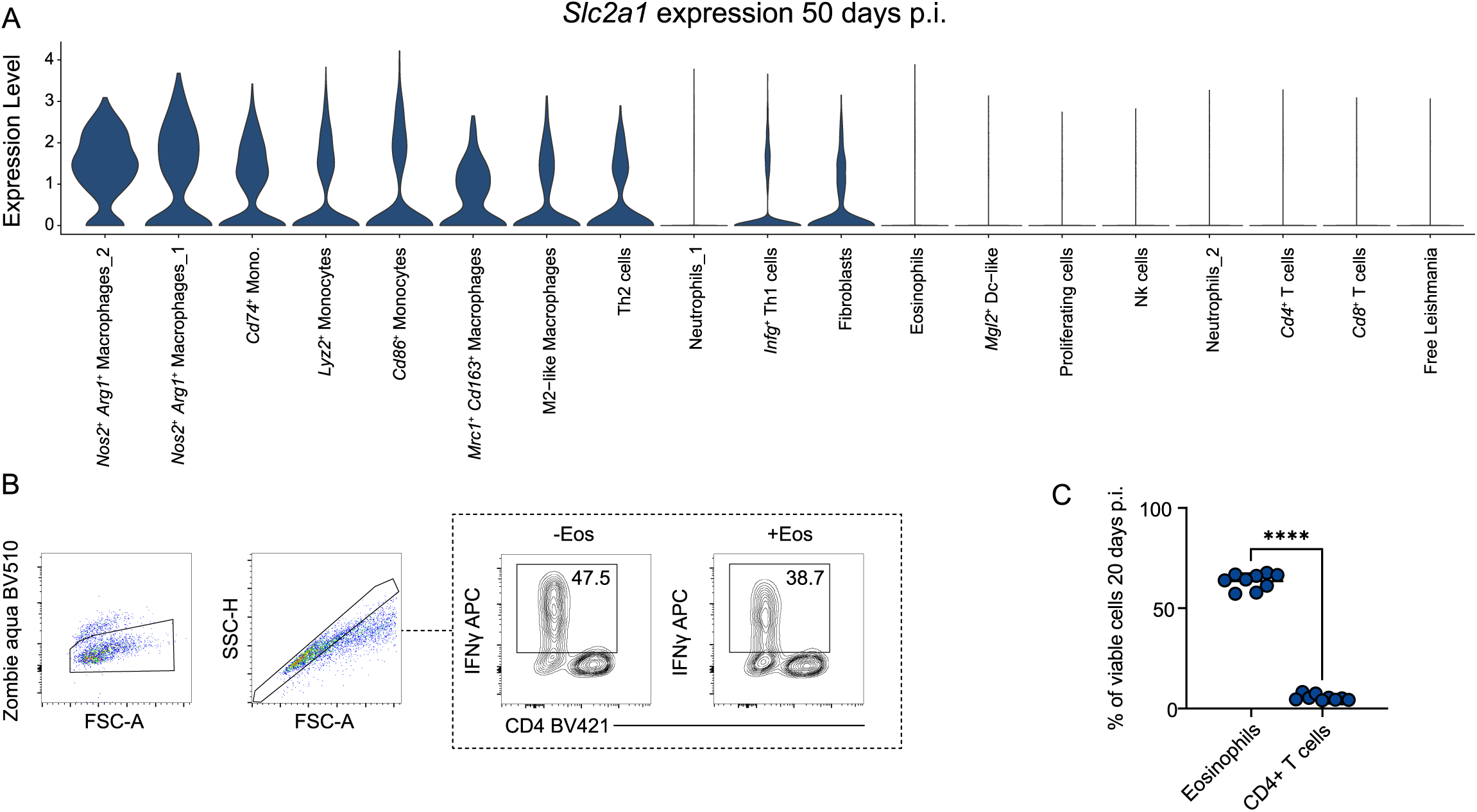
**A** Ranked violin plot of the *Slc2a1* (solute carrier family 2 member 1, glucose transporter 1) expression within all identified clusters at day 50 p.i. (total viable foot skin cells). **B** Exemplary gating strategy for the eosinophils/Th1 co-culture. **C** Flow cytometric quantification of the relative abundance of CD11b^+^ SiglecF^+^ eosinophils and CD3^+^ CD4^+^ T cells 20 days p. i. in *L. mexicana*-induced skin lesions (n = 9, 2 independent experiments). ****p ≤ 0.0001, two-tailed Mann-Whitney U test. Data are mean ± s.d. (error bars fall within the symbols).

## MATERIALS and METHODS

### Mice

All experiments were performed with 6 to 14 week old female mice that were kept under specific pathogen-free conditions with water and food ad libitum. Female mice of a given strain were randomly divided into the different groups. C57BL/6N were obtained from Charles River (Sulzfeld, Germany). IL-4/eGFP 4get reporter mice (Mohrs et al., 2001), eosinophil-deficient ΔdblGATA-1 4get mice (Voehringer et al., 2007; Yu et al., 2002) and ILC2-deficient RorαΔTek mice (Knipfer et al., 2019), all on a C57BL/6 background, have been described before. T. Winkler and S. Zundler (all Erlangen, Germany) kindly supplied Rag1^-/-^ mice on C57BL/6 background (Mombaerts et al., 1992). All animal experiments were approved by the regional animal welfare committee of the governments of Middle or Lower Franconia, respectively (AZ: RUF-55.2.2-2532-2-1461-14). For infection experiments, female knockout or transgenic mice and their age-matched wild type (WT) littermate controls were used. The health status of the mice was closely monitored on a daily basis.

### Parasites and infection

Promastigotes of the *Leishmania mexicana* (*L. mexicana*) strain MNYC/BZ/62/M379 were kindly provided by S. Roberts (Pacific University Oregon, USA) (Roberts et al., 2004). Origin and culture of the parasites have been described elsewhere (Bogdan et al., 2019; Stenger et al., 1996). Mice were infected subcutaneously (s.c.) in the skin of both hind feet with 3 x 10^6^ parasites in 50 µL of PBS. Control mice were left untreated (naïve). The swelling of skin lesions was measured with a metric caliper (Kroeplin) and related to the footpad thickness before infection.

### In vivo treatment

To block the initial upregulation of serum IL-5, *L. mexicana*-infected C57BL6/N mice were injected intraperitoneally with 500 µg of either a blocking anti-mouse IL-5 antibody (InVivoMAb anti-mouse IL-5, BioXCell) or the respective isotype control (InVivoMAb anti-HRP, BioXCell) in 100 µL of PBS. The treatment was administered on days 5 and 12 p.i.

### Quantification of parasite burden by limiting dilution assay

Tissue parasite burden was assessed using limiting dilution analysis as described (Stenger et al., 1996). Organ suspensions were subjected to serial 3-, 5- or 6-fold dilutions (depending on the state of disease progression) in modified Schneideŕs *Drosophila* medium (Leitherer et al., 2017), with 12 replicates per dilution step. Statistical significance was assumed when 95% confidence intervals did not overlap.

### Preparation of single cell suspension from tissue

#### Foot skin

Whole foot skin was cut into small pieces, which were digested for 45 to 60 minutes at 37°C under gentle agitation in RPMI 1640 containing 0.2 mg/mL collagenase P and 0.1 mg/mL DNAse I (both Roche Diagnostics GmbH). Cell suspensions were passed through a 70 µm and 40 µm cell strainer (BD Bioscience), resuspended in 2.5 mL of 40% Percoll preparation (20 mL Percoll [GE Healthcare], 2.2 mL 10×PBS, 27.8 mL HBSS without Ca^2+^ and Mg^2+^ [Sigma-Aldrich]) and layered onto 1.5 mL 60% Percoll (30 mL Percoll, 3.3 mL 10×PBS, 16.7 mL HBSS). After centrifugation for 20 min at 931 x g (room temperature), the cell layer at the interface was collected, washed and resuspended in PBS. In case skin samples were further processed for a scRNA-seq, dead cells were removed using the MACS dead cell removal kit (Miltenyi Biotec) according to the manufacturer’s instructions.

#### Bone marrow (BM)

Femur and tibia were flushed using complete RPMI medium and a 23-gauge needle (BD Biosciences). The content was collected, filtered through a 70 µm cell strainer and red blood cells were lysed using Red Blood Cell Lysis Buffer (Sigma-Aldrich) according to the manufacturer’s protocol. In case BM samples were further processed for a scRNA-Seq, a density gradient centrifugation was then conducted using Histopaque 1077/1119 (Sigma-Aldrich), following the manufacturers protocol. Afterwards, the granulocyte layer was carefully transferred and washed with PBS. The viability was increased by removing dead cells using the MACS dead cell removal kit according to the manufacturer’s instructions *Blood:* Following euthanasia of the mice, a midline laparotomy was performed to expose the abdominal cavity, and the vena cava was carefully isolated and accessed using a 26G needle attached to an Omnifix-F Luer Solo syringe (Braun). Blood was drawn from the vena cava and transferred into a 1.5 mL Eppendorf tube (Eppendorf) for serum analysis or into EDTA-coated tubes (Micro tube 1.3 mL, Sarstedt). In case blood samples were further processed for a scRNA-seq, a density gradient centrifugation was then conducted using Histopaque 1077/1119 (Sigma-Aldrich) following the manufactureŕs protocol. Afterwards, the granulocyte layer was carefully transferred and washed with PBS. Red blood cells were lysed twice in ice-cold water for 10 seconds. The viability was increased by removing dead cells using the MACS dead cell removal kit according to the manufacturer’s instructions.

### Serum analysis

The collected blood samples were then centrifuged at 4°C, 1500 × g for 10 minutes and the upper serum layer was carefully pipetted into a new 1.5 mL Eppendorf tubes.

#### ELISA

The ELISA MAX™ Standard Set Mouse IL-5 (BioLegend) was utilized according to the manufacturer’s instructions, except that serum samples were incubated overnight at 4°C to enhance the detectability of the generally low serum IL-5 concentrations.

#### Legendplex

The Legendplex Mouse Th Cytokine Panel V03 (BioLegend, San Diego) was used according to the manufacturer’s protocol. Serum samples were diluted 2-fold using the provided assay buffer.

### Flow cytometry

Cells resuspended in PBS were stained with the Zombie Aqua^TM^ Fixable Viability Kit (BioLegend) according to the manufacturer’s protocol and washed with PBS/ 1% FCS/ 2 mM EDTA. After incubation with TruStain fcX α-mouse CD16/32 blocking antibody (BioLegend), staining of different surface markers (see Table 1) was performed for 20 min at 4°C followed by a washing step with PBS/1% FCS. In the case of anti-CD34, the antibody incubation was extended to 90 min at room temperature (RT).

For intracellular cytokine, staining cells were fixed with BD Cytofix Cytoperm (BD Biosciences) for 20 min at 4°C, washed twice with 1x saponin buffer (0.5% (w/v) saponin [Carl Roth], 2 mM EDTA, 2% FCS, in PBS) and stained for IFNγ (see Table 1) overnight at 4°C in 1× saponin buffer. Samples were measured using a BD LSRFortessa™ flow cytometer equipped with an ultra-violet (355 nm), blue (488 nm), yellow-green (561 nm), red (640 nm) and violet (605 nm) laser. Flow cytometry data analysis was performed using FlowJo software (v 10.6.1, Becton Dickinson). Cell counts, relative cell frequencies or MFI were used to generate graphical plots in GraphPad Prism (v.9, GraphPad).

**Table 1:** Antibodies used for flow cytometric analysis.

| Antibody | Source | Order No. | Batch, Clone | Fluorochrome Conjugation | Dilution |
| --- | --- | --- | --- | --- | --- |
| Rat anti-mouse CD3 | BD | 740268 | 17A2 | BUV395 | 1:100 |
| Rat anti-mouse CD4 | eBioscience | 25-0042 | GK1.5 | PE-Cy7 | 1:100 |
| Rat anti-mouse CD4 | BioLegend | 100443 | GK1.5 | BV421 | 1:100 |
| Rat anti-mouse CD8a | eBioscience | 17-0081 | 53-6.7 | APC eF780 | 1:100 |
| Rat anti-mouse CD11b | BD | 612800 | M1/70 | BUV737 | 1:200 |
| Rat anti-mouse CD34 | BD | 551387 | RAM34 | PE | 1:50 |
| Rat anti-mouse CD44 | BD | 612799 | IM7 | BUV737 | 1:800 |
| Rat anti-mouse CD45 | BioLegend | 103108 | 30-F11 | FITC | 1:100 |
| Rat anti-mouse CD125 | BioLegend | 153406 | DIH37 | APC | 1:100 |
| Rat anti-mouse CD163 | Thermo Fisher | 12-1631-82 | TNKUPJ | PE | 1:100 |
| Rat anti-mouse CD206 | BioLegend | 141717 | C068C2 | BV421 | 1:100 |
| Rat anti-mouse IFNγ | eBioscience | 505810 | XMG1.2 | APC | 1:100 |
| Rat anti-mouse Ly6G | eBioscience | 47-9668-82 | 1A8 | APC eF780 | 1:100 |
| Rat anti-mouse Ly6G | BioLegend | 127606 | 1A8 | FITC | 1:100 |
| Rat anti-mouse Nk1.1 | BioLegend | 108741 | PK136 | PE-Cy7 | 1:100 |
| Rat anti-mouse SiglecF | eBioscience | 61-1702-82 | 1RNM44N | PE-CF594 | 1:400 |

### 2-NBDG Assay

Single cell suspensions from C57BL/6N skin lesions at day 20 p.i. were prepared as described above. Cells were transferred into 96-well V-bottom plate and allowed to rest for 1h at 37°C, 5% CO_2_/95% humidified air. Afterwards cells were washed in RPMI 1640 (Life Technologies, without Glucose) and resuspended in RPMI 1640 (without Glucose, 100 µM 2-NBDG (Life Technologies), 10% dialyzed FCS). Cells were allowed to rest for 45 min at 37°C and 5% CO_2_/95% humidified air. Next, cells were stained for flow cytometric analysis as described above.

### Sorting of eosinophils

Skin cells from C57BL/6N mice were isolated at day 20 p.i. and stained with anti-Ly6G-FITC (BioLegend, 127606) and anti-SiglecF-PE-CF594 (eBioscience, 61-1702-82). Eosinophils (SiglecF^+^ Ly6G^-^ SSC^High^) were sorted in PBS/ 1% FCS/ 5 mM EDTA using a BioRad S3 Cell Sorter (Bio-Rad).

### Immunofluorescence staining

1×10^5^ eosinophils sorted from the site of *L. mexicana* infection were placed on an ImmunoSelect Adhesion Slide (Squarix). Cells were fixed with 4% PFA, permeabilized with PBS (0.5% Triton-X) and stained overnight at 4°C using the serum of a CL patient (SE6209/13) with high antibody titer against *L. mexicana*. The secondary donkey-anti-human A647 (H+L(ab)2 fragment, Dianova) was added for 1h at RT. Finally, slides were mounted with DAPI Fluoromount G (Southern Biotec). Slides were kept overnight at R, and images were taken using a Keyence fluorescence microscope (Keyence, BZ-X800/ BZ-X810) with a 40x or 100x objective.

### Co-culture of sorted eosinophils and generated Th1 cells

Naïve CD4^+^CD62L^+^ T cells, purified from the spleens of uninfected C57BL/6N mice using the naïve CD4^+^ T cell isolation kit (Miltenyi Biotec), were stimulated for 3 days with immobilized anti-CD3 (clone 145–2C11, BioXCell; culture wells coated with 5 μg/mL) in the presence of Th1-skewed conditions, i.e. with addition of IL-12 (R&D System, 419-ML, 5 ng/µL), anti-IL-4 (R&D System, 404-ML-050/CF, 1 ng/µL) and IL-2 (R&D System, 402-ML-100/CF, 20 ng/µL). Afterwards, cells were split into fresh uncoated wells and expanded with IL-2 (20 ng/µL) for 2 days at 37°C and 5% CO_2_/95% humidified air. 1×10^4^ Th1 cells were co-cultivated in a 96 well plate (Thermo Fisher Scientific) in a 1:1 ratio with sorted eosinophils using RPMI 1640 (20 ng/µL IL-2, 10 ng/µL IL-5, 20% dialyzed FCS) overnight at 37°C and 5% CO_2_/95% humidified air. Co-cultures were stimulated with 50 ng/mL phorbol-12-myristate-13-acetate (PMA) and 750 ng/mL ionomycin (both Sigma-Aldrich) for 4h in the presence of 2 µM monensin (Biolegend) at 37°C and 5% CO_2/_/95% humidified air. Afterwards, viability, cell surface antigen and intracellular cytokine staining was performed as described above.

### Quantitative real-time PCR

Total RNA was extracted from homogenized tissue or cell culture and reverse transcribed as described previously(Schleicher et al., 2016). The following gene-specific assays (TaqMan Gene Expression Assays-on-Demand; Thermo Fisher Scientific) were used for quantitative real-time PCR:

**Table 2:**
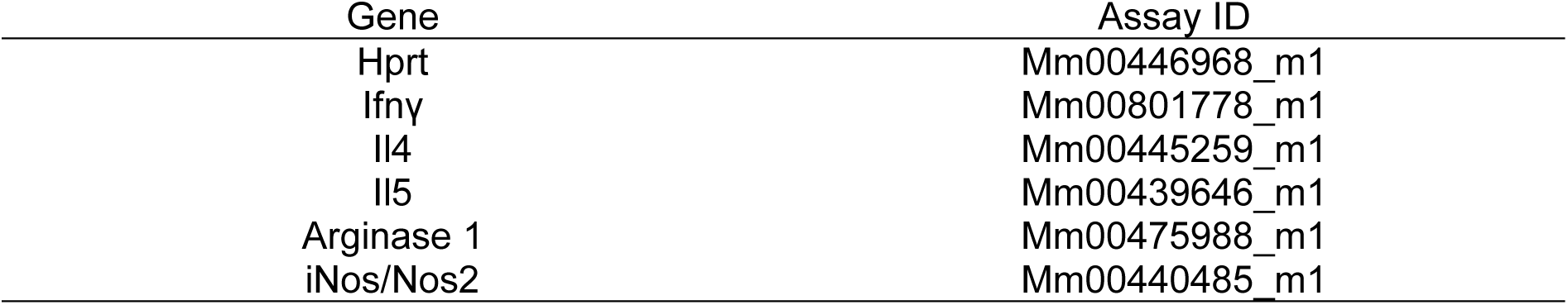
Gene-specific assays used for qRT-PCR.

The mRNA levels were calculated using the following formula:

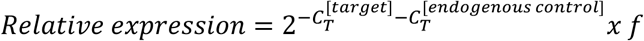

with *C_T_* denoting the cycle threshold and *f* = 10^4^ as an arbitrary factor. In some experiments, relative expression was calibrated to controls, as indicated in the figure legends.

### Transcriptomics

#### Single-cell capture and library preparation

Two independent scRNA-seq experiments were conducted using BD Rhapsody (BD Bioscience). The first experiment included bone marrow, blood and skin cells collected at day 14 p.i. (D14). The second experiment focused on total skin cells collected at day 50 p.i. and compared dblGATA-1 4get C57BL/6 mice and 4get C57BL/6 mice (D50). Tissue processing was performed as described above. Twelve mice were used for D14 and seven mice per group for D50.

Discrimination between viable and dead cells was achieved by staining with 5 μM calcein AM and 0.3 μM DRAQ 7, as described in the Single-Cell Capture and cDNA Synthesis Protocol for BD Rhapsody (BD Bioscience). Samples which were subsequently pooled and labelled with sample tags (BD Bioscience, BD Mouse Single-Cell Multiplexing Set, 626545, rat anti-mouse MHC-H2 class I [clone M1/42]) according to the manufacturer’s protocol. In brief, up to 1 million cells were resuspended in a total of 180 μL freshly prepared FACS Buffer (PBS, 1% FSC, 2mM EDTA). Sample tag tubes from the BD Mouse Single-Cell Multiplexing Kit were briefly centrifuged. 20 μL from the sample tag tube were transferred into the respective sample and incubated at room temperature for 20 minutes in the dark. Each labelled cell suspension was transferred to a 5 mL polystyrene (Falcon) tube and 2 mL FACS buffer were added. Each sample was centrifuged at 1,400 rpm for 6 minutes at 4°C. Supernatant was carefully removed and 2 mL FACS buffer were added, which was repeated twice. Total amount of cells was determined as described above. Cell loading in the BD Rhapsody 8-Lane Cartridge (BD Bioscience) and single-cell capture with the BD Rhapsody HT Single-Cell Analysis System was performed as mentioned in the manufacturer’s instructions. Afterwards, single-cell whole transcriptome mRNA and sample tag libraries were generated according to the manufacturer’s instructions. The final libraries were quantified using a Qubit Fluorometer (Thermo Fisher Scientific) with the Qubit dsDNA HS Assay Kit (Thermo Fisher Scientific). Library size distribution was measured with the Agilent High Sensitivity D5000 assay (Agilent) on a TapeStation 4200 (Agilent) system. Sequencing was performed on a NovaSeq 6000.

#### Data pre-processing and normalization

Paired-end scRNA-seq FASTQ files were processed on the Seven Bridges Genomics platform with default parameters. A customized reference database was created using the *Leishmania mexicana* genome (Genome assembly ASM23466v4, Wellcome Trust Sanger Institute) and the *Mus musculus* genome (GRCm38, modified to meet Cell Ranger requirements). This reference genome was used for D50, whereas D14 was solely aligned to the *Mus musculus* genome. Downstream analysis was conducted in R (v.4.4.0) using the Seurat package v.5.1.0(Hao et al., 2024). All Seurat objects (one for each multiplexed sample) were merged if necessary and subjected to the same quality filtering. Cells with fewer than 200 or more than 6,000 detected genes were excluded from the analysis. Following log normalization, the count data were scaled while regressing out mitochondrial reads and principal component analysis (PCA) was performed based on the 2,000 most variable features. To identify and exclude potential doublets, we employed the scDblFinder(Germain et al., 2021) especially for D50. Doublet prediction was performed on the integrated dataset, accounting for batch effects. To enhance efficiency, the prediction process utilized custom cell population annotations. Clustering and UMAP visualization were conducted on the merged dataset using 10 to 30 principal components and a resolution between 0.4 and 1.2 for the shared nearest neighbour clustering algorithm (depending on the respective dataset). The clusters were manually annotated based on marker gene expression. Cell clusters of interest were further divided into subsets and subjected to normalization, scaling and PCA as described above.

#### Differential gene expression analysis and score computation

To extract cluster markers, the FindAllMarkers function was executed with log fold change (logFC) threshold and minimum percentage (min.pct) cut-offs set to 0.25. Top-ranked genes, based on logFC, were extracted for illustration. For differential gene expression analysis, the FindMarkers function was applied with the same logFC threshold and min.pct cut-offs. Genes were subsequently filtered based on Bonferroni-adjusted P-values of less than 0.05. Scores were computed using the AddModuleScore function. The genes used for the glycolysis gene scores and signatures were manually curated from Gene Ontology (GO) ID: 0006006.

#### Eosinophil identification

Eosinophils were identified in silico based on the simultaneous expression of canonical marker genes, including *Il5ra*, *Ccr3*, *Siglecf* and *Prg2*, for both D14 and D50 samples. For the D14 dataset, several clusters exhibiting high expression of neutrophil markers, including but not limited to *Ly6g*, were excluded from the analysis. No eosinophils were identified from the bone marrow (BM) samples.

#### Data integration

Published scRNA-seq data, along with gene scores derived from bulk sequencing datasets, were utilized to integrate and contextualize the subsets of the eosinophil cluster (see Table 3). To integrate human scRNA-seq(Cui et al., 2024) data with our mouse data (see Figure 7E), we first subsetted eosinophils based on the expression of *SIGLEC8*. Following this, we converted human genes to their corresponding mouse homologs using the R package biomaRt (v. 3.19(Durinck et al., 2009)) and homologene (v. 1.4.68). Finally, the R package Harmony (v. 1.2.1(Korsunsky et al., 2019)) was employed for accurate integration of the scRNA-Seq data, effectively correcting for batch effects. Gene scores from bulk sequencing were computed using the AddModuleScore function in Seurat. Before conducting the longitudinal integration (see Figure 7G), we removed all *Leishmania* genes sequenced on day 50. Subsequently, we intersected the gene lists from both time points, integrating only the shared genes back into their respective Seurat objects.

**Table 3:** Integrated published datasets.

| Integrated dataset | Source Publication |
| --- | --- |
| Subset of liver eosinophils | (Wang et al., 2022) |
| Top differentially upregulated genes from<br>Clec4a4<sup>+/-</sup> Eos | (Cui et al., 2024) |

### Statistical analysis

Statistical analysis was conducted using either GraphPad Prism (v. 9, GraphPad) or R (v. 4.4.0). Before performing statistical tests, all data sets were tested for Gaussian distribution and outliers were identified using ROUT method. Statistical significance tests were performed as described in each figure legend. P-values ≤ 0.05 were considered as significant. Plots generated in R were visualized using R package ggplot2 (v. 3.5.1).

## Data availability

ScRNA-seq data generated during this study have been deposited at the Gene Expression Omnibus under the accession numbers GSE281802 and GSE281715.

## Supplemental materials

The supplementary figures S1, S2, S3, S4 and S5 are provided as supplementary material.

## Notes

https://www.ncbi.nlm.nih.gov/geo/

### Competing Interest Statement

The authors have declared no competing interest.

